# Probing the role of synaptic adhesion molecule RTN4RL2 in setting up cochlear connectivity

**DOI:** 10.1101/2024.09.16.613011

**Authors:** Nare Karagulyan, Maja Überegger, Yumeng Qi, Norbert Babai, Florian Hofer, Lejo Johnson Chacko, Fangfang Wang, Maria Luque, Rudolf Glueckert, Anneliese Schrott-Fischer, Yunfeng Hua, Tobias Moser, Christine Bandtlow

## Abstract

Sound encoding depends on the precise and reliable neurotransmission at the afferent synapses between the sensory inner hair cells (IHCs) and spiral ganglion neurons (SGNs). The molecular mechanisms contributing to the formation, as well as interplay between the pre- and postsynaptic components remain largely unclear. Here, we tested the role of the synaptic adhesion molecule and Nogo/RTN4 receptor homolog RTN4RL2 (also referred to as NgR2) in the development and function of afferent IHC-SGN synapses. Upon deletion of RTN4RL2 in mice (RTN4RL2^-/-^), presynaptic IHC active zones showed enlarged synaptic ribbons and a depolarized shift in the activation of Ca_V_1.3 Ca^2+^ channels. The postsynaptic densities (PSDs) of SGNs were smaller and deficient of GluA2-4 AMPA receptor subunits despite maintained Gria2 mRNA expression in SGNs. Next to synaptically engaged PSDs we observed “orphan” PSDs located away from IHCs. They likely belong to a subset of SGN peripheral neurites that do not contact the IHCs in RTN4RL2^-/-^ cochleae as found by volume electron microscopy reconstruction of SGN neurites. Auditory brainstem responses of RTN4RL2^-/-^ mice showed increased sound thresholds indicating impaired hearing. Together, these findings suggest that RTN4RL2 contributes to the proper formation and function of auditory afferent synapses and is critical for normal hearing.

## Introduction

Hearing relies upon the correct formation and maturation of cochlear afferent synapses between postsynaptic type I spiral ganglion neurons (SGNs) and presynaptic inner hair cells (IHCs; reviewed in Johnson et al., 2019; Bulankina and Moser, 2012; Appler and Goodrich, 2011). In mice, SGN neurites extend and reach IHCs at late embryonic stages (E16.5; Koundakjian, Appler, and Goodrich, 2007), whereby early synaptic contacts are established between the two cells at E18 (Michanski et al., 2019). Developmental changes and maturation of afferent synapses continue until the onset of hearing (p12), with further refinement occurring till the fourth postnatal week (Wong et al., 2014; Liberman and Liberman, 2016; Michanski et al., 2019). In mature cochleae, each IHC receives a contact from 5-30 SGNs, while each SGN contacts only one active zone (AZ) of one IHC in the majority of cases (Meyer and Moser, 2010; Hua et al., 2021).

To this date, multiple mechanisms have been implicated in SGN neurite guidance and establishment of IHC innervation by SGNs, including signaling via EphrinA5-EphA4, Neuropilin2/Semaphorin3F, Semaphorin5B/PlexinA1, Semaphorin3A (Defourny et al., 2013; Coate et al., 2015; Jung et al., 2019; Cantu-Guerra et al., 2023). Less is known about the molecules governing the transsynaptic organization at IHC afferent synapses. Similar to the central conventional synapses, synaptic adhesion proteins such as neurexins and neuroligins have been suggested to play a role in pre- and postsynaptic assemblies (Ramirez et al., 2022; Jukic et al., 2024). Furthermore, the Ca_V_1.3 extracellular auxiliary subunit Ca_V_α_2_δ_2_ was indicated to be important for proper alignment of presynaptic IHC AZs and postsynaptic densities (PSDs) of SGNs (Fell et al., 2015).

Whether reticulon 4 receptors (RTN4Rs) contribute to setting up afferent connectivity in the cochlea remained to be investigated. The RTN4 receptor family consists of 3 homologous proteins RTN4R, RTN4RL1 and RTN4RL2. RTN4Rs are leucine-rich repeat (LRR) and glycosylphosphatidylinositol (GPI) anchored cell surface receptors (Figure 1A). Their primary role is thought to limit synaptic plasticity and axonal outgrowth as well as to restrict axonal regeneration after injury (reviewed in Mironova and Giger, 2013). While RTN4R and RTN4RL1 are involved in axonal guidance (Vaccaro et al., 2022), RTN4RL2 has been proposed to be important for innervation of the epidermis by dorsal root ganglion neurons (Bäumer et al., 2014). Furthermore, RTN4Rs are suggested to control the number and the development of synapses (Wills et al., 2012) and to play a role in transsynaptic signaling (Wang et al., 2021). Recent single-cell transcriptomic studies of the cochlea detected the expression of RTN4Rs in SGNs and IHCs (Shrestha et al., 2018; Jean et al., 2023). Here, we investigated the role of RTN4RL2 in the cochlea using previously described RTN4RL2 constitutive knock-out mice (RTN4RL2^-/-^; (Wörter et al., 2009)). We found RTN4RL2 to be expressed in SGNs and to be required for normal hearing: auditory brainstem responses were impaired upon RTN4RL2 deletion. We discovered both pre- and postsynaptic alterations of IHC-SGN synapses in RTN4RL2^-/-^ mice: presynaptic Ca^2+^ channels of IHCs required stronger depolarization to activate and PSDs seemed deficient of GluA2-4 AMPA receptor subunits. Additionally, a subset of type I SGN neurites did not contact IHCs but likely still feature “orphan” PSDs.

**Figure 1.**
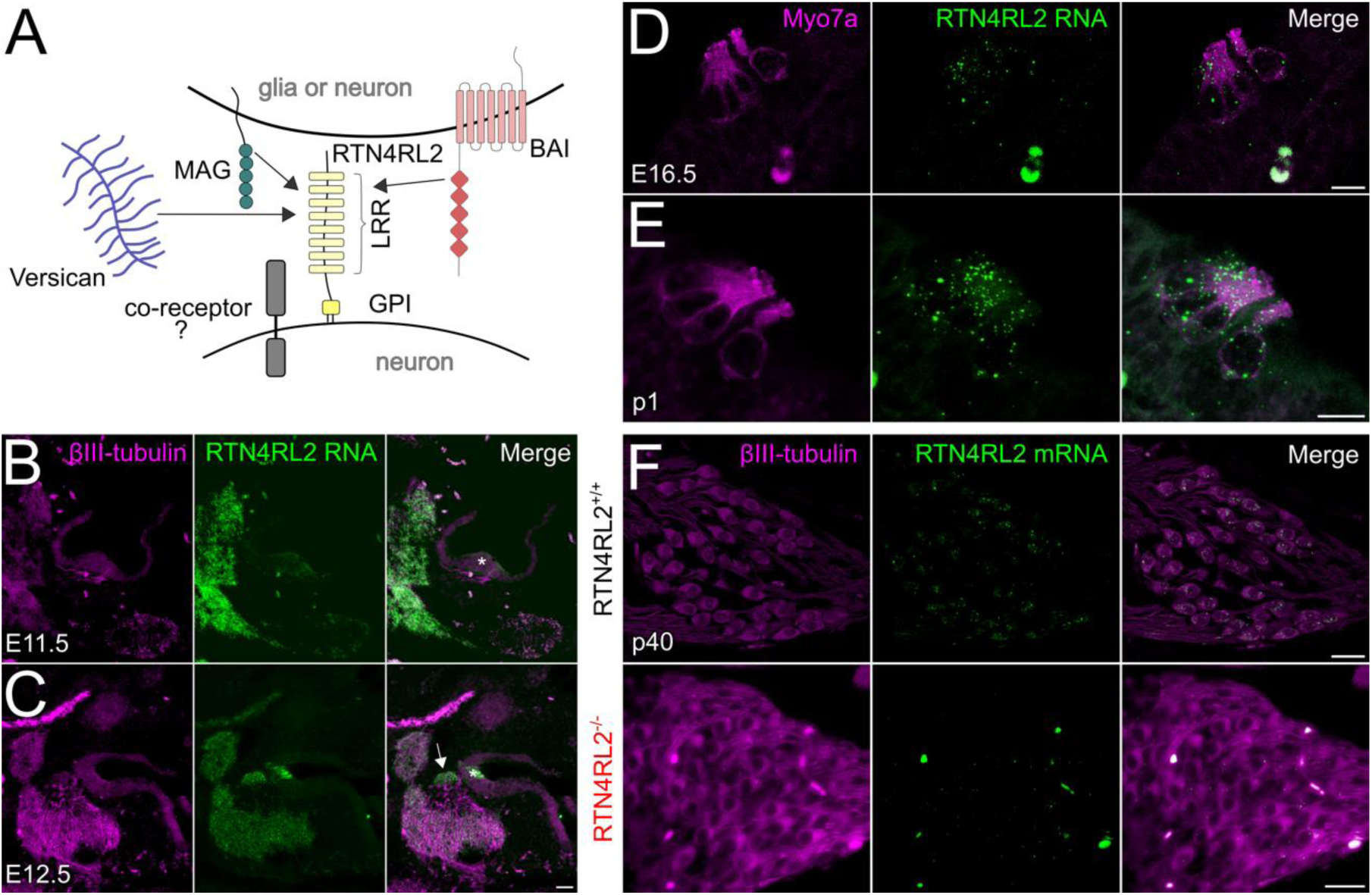
RTN4RL2 mRNA expression in IHCs and SGNs of the mouse cochlea. (**A**) RTN4RL2 is an LRR protein and is anchored to the cell membrane via GPI. In the nervous system RTN4RL2 has been implicated to interact with MAG, versican, BAI (Venkatesh et al., 2005; Bäumer et al., 2014; Wang et al., 2021). (**B, C**) Exemplary images of RNAscope ISH for RTN4RL2 mRNA (green) in the otic vesicles of E11.5 (B) and E12.5 (C) mice. Scale bar = 50 μm. The developing organ of Corti region is marked with an asterisk and the spiral ganglion is indicated with an arrow. (**D, E**) Exemplary images of RTN4RL2 RNA expression in hair cells of E16.5 (D) and p1 (E) mice. Hair cells are visualized with anti-Myo7a stainings. Scale bars = 10 μm. (**F**) Representative images of RNAscope ISH for RTN4RL2 mRNA (green dots) combined with immunostaining for neuron-specific marker βIII-tubulin (grey) in paraffin sections of p40 RTN4RL2^+/+^ and RTN4RL2^-/-^ cochleae. Scale bars = 20 μm.

## Results

### RTN4RL2 is expressed both in hair cells and spiral ganglion neurons

Recent transcriptomic data had detected RTN4RL2 expression both in hair cells and SGNs (Elkon et al., 2015; Liu et al., 2018; Shrestha et al., 2018; Jean et al., 2023). We verified this by performing RNAscope staining in cochleae during inner ear development. We observed RTN4RL2 mRNA expression at E11.5 in the hair cell region and at E12.5 in both the spiral ganglion and hair cells (Figure 1B, C). RTN4RL2 expression was further evident in hair cells at E16.5 and p1 (Figure 1D, E) and was maintained at low levels in spiral ganglia of p40 mice (Figure 1F). RTN4RL2^-/-^ mice were lacking RTN4RL2-specific mRNA puncta (Figure 1F), demonstrating specific detection of RTN4RL2 expression in SGNs.

### RTN4RL2 is important for the correct development of the auditory afferent synapses

To probe the role of RTN4RL2 in the cochlea, we first studied the numbers of cochlear cells in RTN4RL2^-/-^ mice. We did not observe any change in SGN density or counts of inner and outer hair cells at p15, 1-month- and 2-month-old mice (Figure 2-figure supplement 1). Given that RTN4RL2 has been implicated in synapse formation and development (Wills et al., 2012; Borrie et al., 2014; Wang et al., 2021) we immunolabeled IHC afferent synapses for presynaptic RIBEYE/Ctbp2 and postsynaptic Homer1 in mice at the age of 3 weeks. While we did not detect any change in the number of synaptic ribbons in RTN4RL2^-/-^ IHCs, ribbon volumes were bigger (Figure 2C, D; Figure 2-figure supplement 2). Interestingly, in addition to the Homer1 positive puncta juxtaposing presynaptic ribbons, we observed additional Homer1 patches, which appeared to be away from IHCs, potentially marking “orphan” PSDs (Figure 2A, B). Moreover, the Homer1 puncta juxtaposing IHC AZs were significantly smaller in RTN4RL2^-/-^ mice compared to the control IHCs (Figure 2E). While we found the percentage of presynaptic ribbons juxtaposing Homer1 immunofluorescent puncta to be decreased by approximately 7% in RTN4RL2^-/-^ mice (Figure 2F), we cannot exclude that our immunolabeling protocol lacked the sensitivity to detect smaller synaptically engaged PSDs. Out of the four pore-forming AMPA receptor subunits (GluA1-4), mature SGNs express GluA2-4 (Niedzielski and Wenthold, 1995; Matsubara et al., 1996). Our immunolabeling of anti-GluA2/3 revealed a severe reduction in the number of GluA2/3 positive puncta in RTN4RL2^-/-^, indicating possible lack or decrease of GluA2/3 (Figure 3A, B). Interestingly, the remaining GluA2/3 patches did not properly juxtapose presynaptic ribbons (Figure 3C). Despite this, the expression of Gria2 mRNA appeared to be maintained in SGNs, as indicated by RNAscope (Figure 3D). We then checked whether the potential lack of GluA2/3 was accompanied by an increase in GluA4 signal at the PSDs of RTN4RL2^-/-^ SGNs, as was previously shown in GluA3 KO mice (Rutherford et al., 2023). Surprisingly, we did not detect a significant GluA4 signal compared to the background in RTN4RL2^-/-^ cochleae (Figure 3-figure supplement 1). To check if the “orphan” PSDs away from IHCs were possibly erroneously engaged with efferent presynaptic terminals we stained the latter for synapsin, which is lacking from IHCs, but did not observe any obvious juxtaposition between the “orphan” Homer1 and synapsin puncta (Figure 2-figure supplement 3).

**Figure 2.**
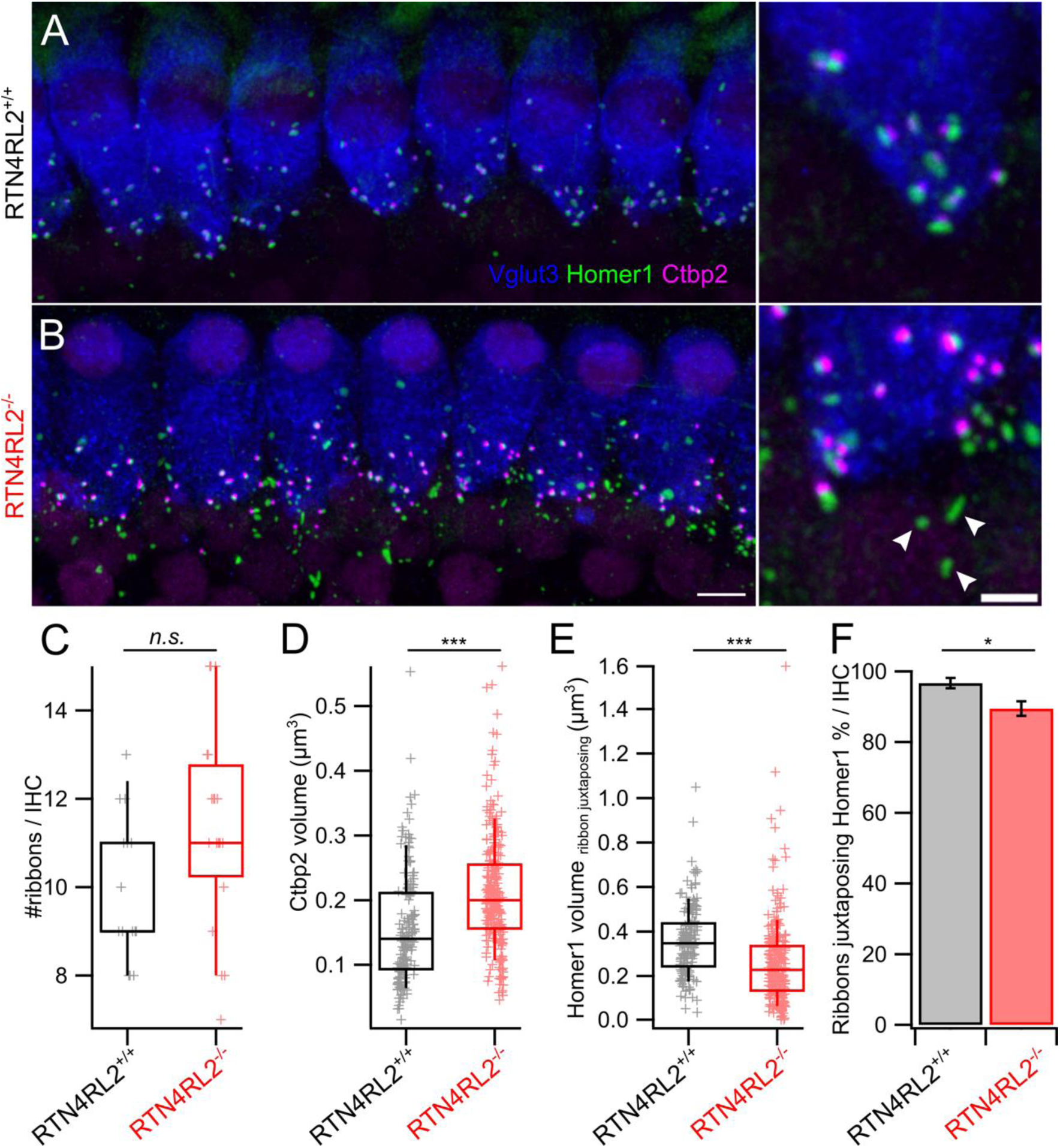
Pre- and postsynaptic changes at IHCs of RTN4RL2^-/-^ mice. (**A, B**) Maximum intensity projections of representative confocal stacks of IHCs from apical cochlear region of 3-week-old RTN4RL2^+/+^ (A) and RTN4RL2^-/-^ (B) mice immunolabeled against Vglut3, Homer1, and Ctbp2. Scale bar = 5 μm. Images on the right-hand side are zoomed into the synaptic regions. Scale bar = 2 μm. Some of the putative “orphan” PSDs are marked with the white arrowheads. (**C**) Number of Ctbp2 positive puncta is not changed in IHCs of RTN4RL2^-/-^ mice (RTN4RL2^+/+^: 9.9 ± 0.42, SD = 1.62, n = 15, N = 2 vs RTN4RL2^-/-^: 11.4 ± 0.5, SD = 2.25, n = 20, N = 3; p = 0.06, Mann-Whitney-Wilcoxon test). (**D**) Ribbon volumes are enlarged in RTN4RL2^-/-^ IHCs (RTN4RL2^+/+^: 0.16 ± 0.007 μm^3^, SD = 0.09 μm^3^, n = 165, N = 2 vs RTN4RL2^-/-^: 0.21 ± 0.005 μm^3^, SD = 0.09 μm^3^, n = 259, N = 3; p < 0.001, Mann-Whitney-Wilcoxon test). (**E**) Homer1 patches which are juxtaposing presynaptic ribbons show decreased volumes in RTN4RL2^-/-^ IHCs (RTN4RL2^+/+^: 0.36 ± 0.01 μm^3^, SD = 0.16 μm^3^, n = 160, N = 2 vs RTN4RL2^-/-^: 0.26 ± 0.01 μm^3^, SD = 0.19 μm^3^, n = 249, N = 3; p < 0.001, Mann-Whitney-Wilcoxon test). (**F**) Percentage of Ctbp2 puncta juxtaposing Homer1 is slightly decreased in RTN4RL2^-/-^ mice (RTN4RL2^+/+^: 96.7 ± 1.45 %, SD = 5.44 %, n = 14, N = 2 vs RTN4RL2^-/-^: 89.5 ± 2.05 %, SD = 9.18 %, n = 20, N = 3; p = 0.03, Mann-Whitney-Wilcoxon test). Data is presented as mean ± SEM. Box-whisker plots show the median, 25/75 percentiles (box), and 10/90 percentiles (whiskers). Individual data points are overlaid.

**Figure 3.**
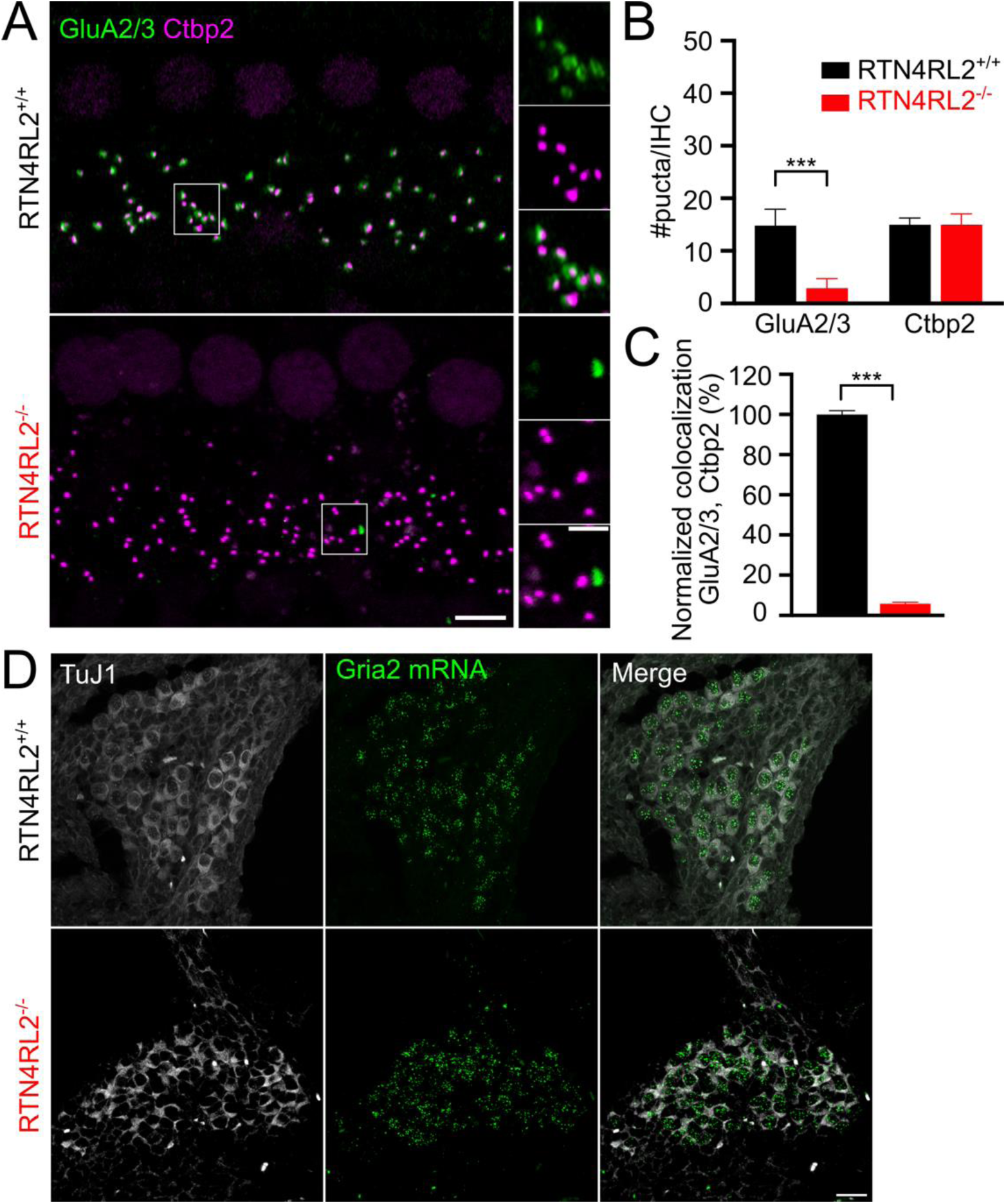
Reduced GluA2/3 signal juxtaposing presynaptic ribbons in IHCs of RTN4RL2^-/-^ mice. (**A**) Maximum intensity projections of representative IHC regions from 1- to 1.5-month-old RTN4RL2^+/+^ (top) and RTN4RL2^-/-^ (bottom) mouse cochleae immunolabeled against Ctbp2/Ribeye (ribbons) and GluA2/3 (AMPA receptors of PSD). Scale bar = 5 μm. The zoom-in regions marked with the white rectagles are presented on the right-hand side. Scale bar = 2 μm. (**B**) The number of the GluA2/3 positive puncta is drastically reduced in RTN4RL2^-/-^ mice despite the maintained number of presynaptic ribbons (p < 0.001, Mann-Whitney-Wilcoxon test). (**C**) Disrupted colocalization of Ctbp2/Ribeye and GluA2 immunofluorescence puncta at IHCs of RTN4RL2^-/-^ mice (p < 0.001, Mann-Whitney-Wilcoxon test). N = 6 animals/genotype. (**D**) Representative images of RNAscope ISH from p4 mice show maintained expression of Gria2 (red dots) in the SGN somata of RTN4RL2^-/-^ mice. Scale bar = 20 μm.

### Additional non-synaptic neurites in the cochlea of RTN4RL2^-/-^ mice

To further examine the afferent cochlear connectivity, we utilized serial block-face scanning electron microscopy (SBEM) to 3D reconstruct the apical cochlea segments from two RTN4RL2^-/-^ and one littermate RTN4RL2^+/+^ control mice at p36 (Figure 4A). As recently demonstrated (Hua et al., 2021; Lu et al., 2024), the spatial resolution of SBEM allows reliable ribbon synapse identification and neurite reconstruction (Figure 4B and C). In these three datasets, we have traced all neural fibers from the habenula perforata (three in each dataset). This resulted in 133 fibers, of which 115 were classified as peripheral neurites of type I SGN based on their radial calibers and extensive myelination after entering the habenula perforata (Figure 4D). We observed a substantial number of neurites that did not engage the IHCs (non-synaptic neurites) in the RTN4RL2^-/-^ organs of Corti (23 out of 83) in contrast to the wild-type control (3 out of 32, Figure 4E). Although most non-synaptic neurites were found in the inner spiral bundle, they failed to reach IHC basal lateral poles to form contacts (17 out of 23 in RTN4RL2^-/-^ animals). This result might provide a plausible explanation for the “orphan” Homer1 puncta shown by immunohistochemistry in RTN4RL2^-/-^ organs of Corti. Note that the presence of non-synaptic neurites in the RTN4RL2^-/-^ animal was not associated with a profound reduction in ribbon synapses (Figure 2C, Figure 2-figure supplement 2B, Figure 4- figure supplement 1A), suggesting a different scenario than deafferentation due to excitotoxic synaptopathy as recently reported (Moverman et al., 2023). Nevertheless, nearly all traced radial fibers were found predominantly unbranched in both the RTN4RL2^-/-^ (97%) and the wild-type control mice (94%; Figure 4F).

**Figure 4.**
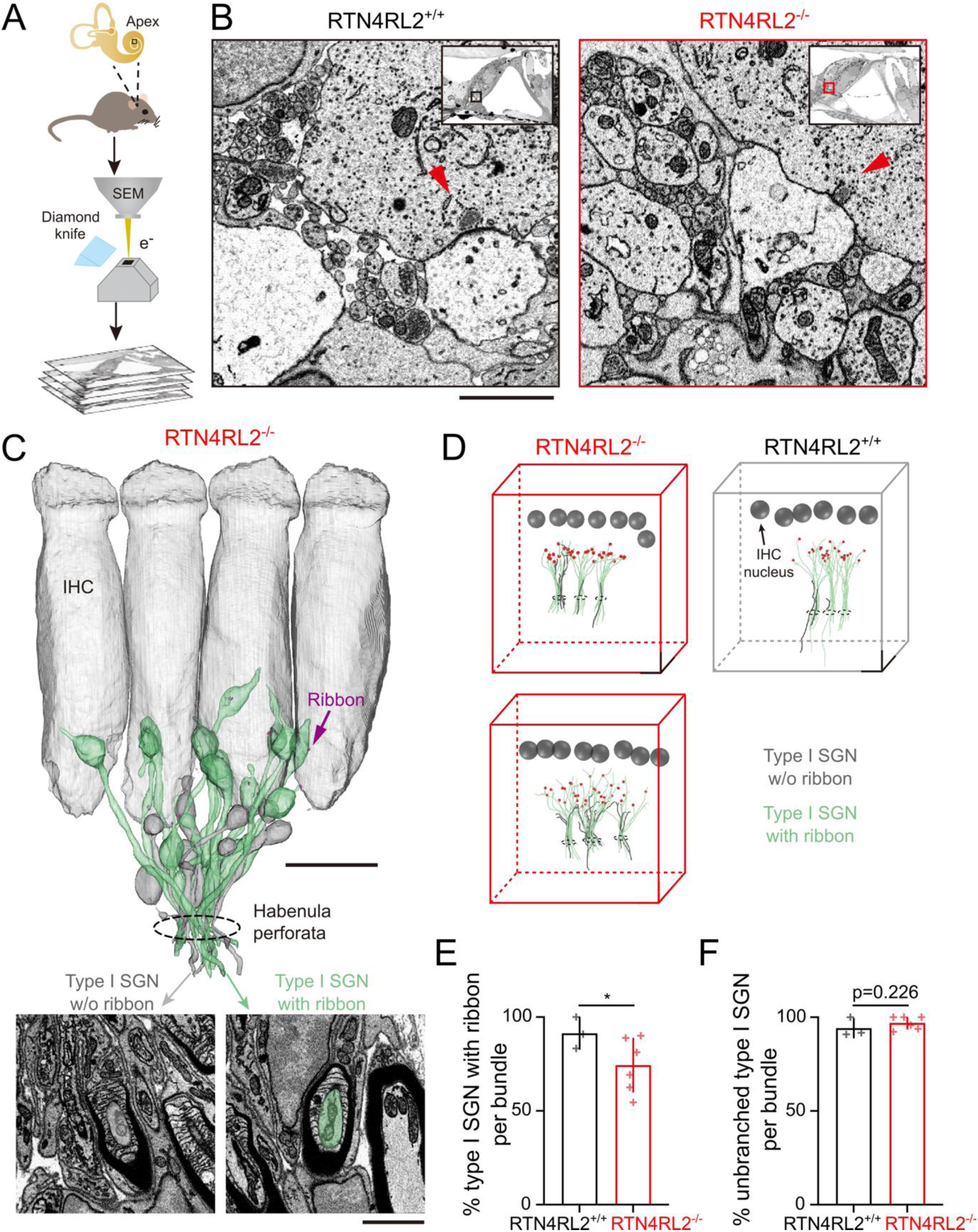
Additional non-synaptically engaged SGN neurites in the cochlea of RTN4RL2^-/-^ mice. (**A**) Workflow of SBEM imaging at the mouse apical cochlear region. (**B**) Example images of neurites beneath IHCs from the RTN4RL2^+/+^ (left) and RTN4RL2^-/-^ (right) mice. Synaptic ribbons are indicated with red arrows. The regions of interest were magnified from single sections of SBEM datasets (insets). Scale bar = 2 μm. (**C**) 3D rendering of afferent fiber reconstruction with ribbons (red), showing both synaptic (green, type I SGN with ribbon) and non-synaptic (grey, type I SGN without ribbon) populations in the RTN4RL2^-/-^ mouse. Scale bar = 10 μm. Inset: EM images show the SGN segments on which myelin sheaths start to form. Scale bar 2 = μm. (**D**) Display of classified radial fibers in the RTN4RL2^+/+^ (right) and RTN4RL2^- /-^ (left and bottom) animals. All fibers were traced from the habenula perforata (cycles) before classification to avoid bias to terminal types. Scale bar = 10 μm. (**E**) Percentage of radial fibers with ribbon per bundle (RTN4RL2^+/+^: 91.41 ± 8.35 %, n = 3 bundles, N = 1 vs RTN4RL2^-/-^: 74.40 ± 14.58%, n = 6 bundles, N = 2; p = 0.032, unpaired t-test). (**F**) Percentage of unbranched radial fibers per bundle (RTN4RL2^+/+^: 94.19 ± 5.04 %, n = 3 bundles, N = 1 vs RTN4RL2^-/-^: 96.98 ± 3.49 %, n = 6 bundles, N = 2; p = 0.226, unpaired t-test).

### Deletion of RTN4RL2 results in depolarized shift of the Ca^2+^ channel activation in IHCs

Next, we tested for potential effects of RTN4RL2 deletion on the presynaptic IHC function. We performed whole-cell patch-clamp recordings from IHCs of p21-29 RTN4RL2^+/+^ and RTN4RL2^-/-^ mice. First, we recorded voltage-gated Ca^2+^ currents by applying step depolarizations with 5 mV increment in ruptured patch-clamp configuration (Figure 5A, see Materials and Methods). We did not observe any change in the maximal Ca^2+^ current amplitude (Figure 5B, C), yet noticed small (∼ +3 mV) but consistent and statistically significant depolarized shift of the voltage of half-maximal Ca^2+^ channel activation (V_half_; Figure 5D, E). The voltage sensitivity of the channels was not changed in RTN4RL2^-/-^ IHCs (Figure 5F). Next, we probed Ca^2+^ influx triggered exocytosis from IHCs by applying depolarizations of varying durations to voltages saturating Ca^2+^ influx in IHCs of both genotypes (-17 mV) and recording exocytic membrane capacitance changes (ΔC_m_) in the perforated patch-clamp configuration (Figure 5G). Both fast (up to 20 ms depolarization) and sustained components of exocytosis remained intact in the mutant, indicating unaffected readily releasable pool and replenishment of the vesicles in RTN4RL2^-/-^ IHCs (Figure 5H).

**Figure 5.**
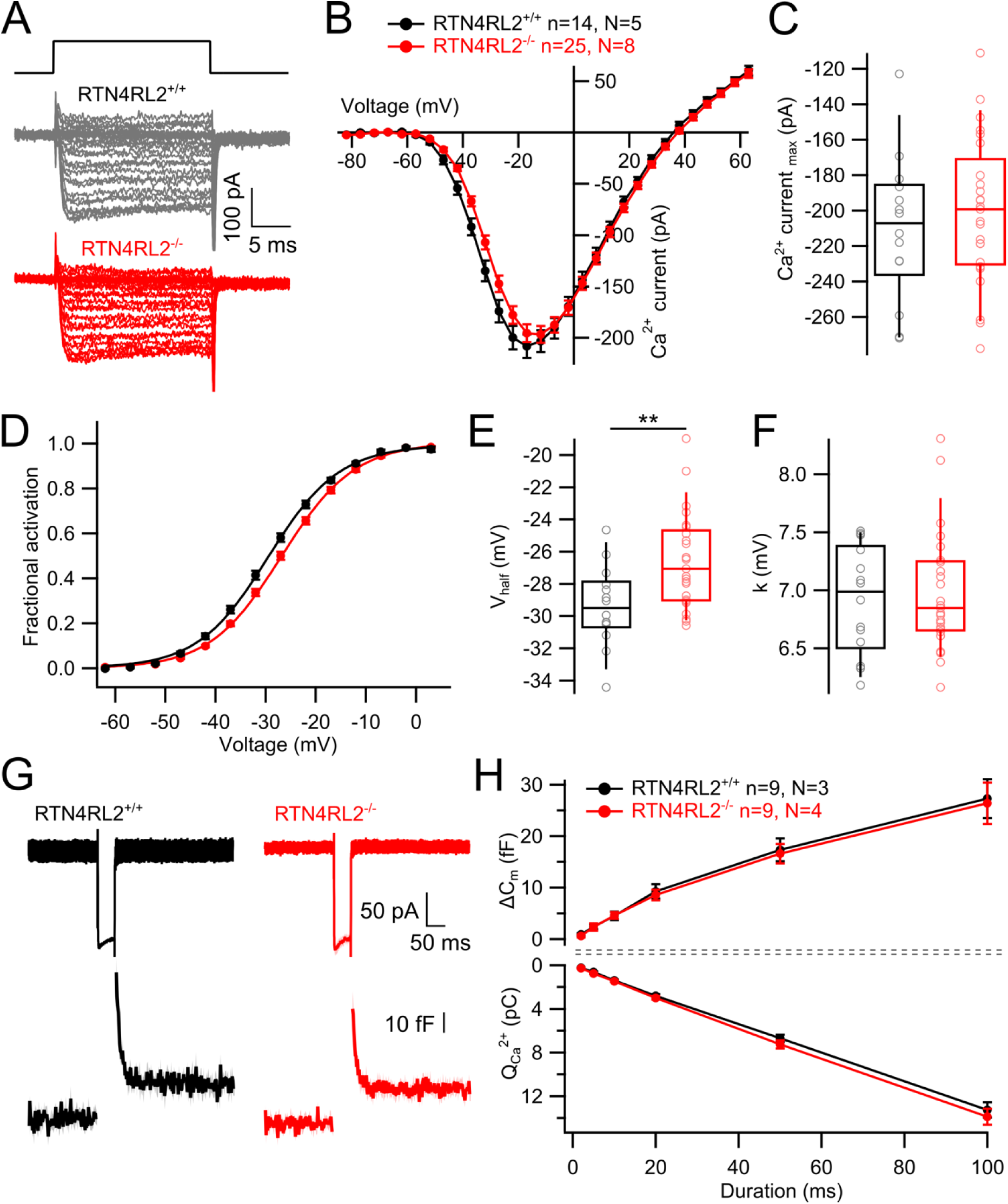
Shifted operation range of Ca^2+^ channels but intact exocytosis in IHCs of RTN4RL2^-^ **^/-^ mice.** (**A**) Representative current traces from IHCs of RTN4RL2^+/+^ (top, black) and RTN4RL2^+/+^ (bottom, red) evoked by step depolarizations. (**B**) Average Ca^2+^ current-voltage relationships (IV curves) in RTN4RL2^+/+^ and RTN4RL2^-/-^ IHCs. (**C**) Maximal Ca^2+^ current amplitude is not changed in RTN4RL2^-/-^ IHCs (RTN4RL2^+/+^: 210 ± 10.9 pA, SD = 40.9 pA, n = 14, N = 5 vs RTN4RL2^-/-^: -200 ± 8.19 pA, SD = 40.9 pA, n = 25, N = 8; p = 0.47, Student’s t-test). (**D**) Fractional activation curves of Ca^2+^ channels calculated from IVs show depolarized shift in channel activation in RTN4RL2^-/-^ IHCs. (**E**) Voltage of half maximal activation obtained from Boltzmann fit of the curves from (D) is more positive in RTN4RL2^-/-^ IHCs (RTN4RL2^+/+^: -29.4 ± 0.66 mV, SD = 2.48 mV, n = 14, N = 5 vs RTN4RL2^-/-^: -26.6 ± 0.59 mV, SD = 2.94 mV, n = 25, N = 8; p = 0.004, Student’s t-test). (**F**) Voltage sensitivity (k) is not changed in RTN4RL2^-/-^ IHCs (RTN4RL2^+/+^: 6.92 ± 0.12 mV, SD = 0.46 mV, n = 14, N = 5 vs RTN4RL2^-/-^: 6.98 ± 0.1 mV, SD = 0.51 mV, n = 25, N = 8; p = 0.93, Mann-Whitney-Wilcoxon test). (**G**) Average current traces evoked by 50 ms depolarization to -17 mV (top row) and resulting capacitance response (bottom row) from RTN4RL2^+/+^ (left, black) and RTN4RL2^+/+^ (right, red) IHCs. Shaded areas represent ± SEM. (**H**) Exocytic capacitance change (ΔC_m_, top) and corresponding Ca^2+^ charge (Q_Ca_^2+^, bottom) evoked by depolarizations (to -17 mV) of various durations (2, 5, 10, 20, 50, 100 ms). Box-whisker plots show the median, 25/75 percentiles (box) and 10/90 percentiles (whiskers). Individual data points are overlaid.

We then turned to study single AZ function by combining whole cell ruptured patch-clamp with spinning disc confocal imaging of presynaptic Ca^2+^ influx, as described previously (Ohn et al., 2016). TAMRA conjugated Ctbp2 binding dimeric peptide and the low affinity Ca^2+^ indicator Fluo4-FF (k_D_: 10 µM) were loaded into the cell via the patch pipette and Ca^2+^ influx at single AZs was visualized in the form of hotspots by applying voltage ramp depolarizations to the IHCs and simultaneously imaging Fluo4-FF fluorescence near labeled ribbons (Figure 6A, see Materials and methods). Maximal Ca^2+^ influx at single AZs tended to be higher in IHCs of RTN4RL2^-/-^ mice but the difference did not reach statistical significance (Figure 6B, B^i^). The V_half_ of Ca^2+^ channel clusters at single AZs showed depolarized shift (∼ +5 mV), which was statistically significant despite the large variability of V_half_ across different AZs (Figure 6C, C^i^). Furthermore, we did not observe any mismatch in Ctbp2 positive puncta and evoked Ca^2+^ hotspots, indicating the correct localization of Ca^2+^ channels at the AZs. We verified this by performing immunostainings of Ca_V_1.3, Ctbp2 and Bassoon and did not detect any apparent misalignment of these protein clusters (Figure 6D). In summary, deletion of RTN4RL2 causes a depolarized shift of the activation of presynaptic Ca^2+^ influx in IHCs as demonstrated at the single AZ and whole cell levels. This is expected to elevate the sound pressure levels required to reach a comparable activity of synaptic transmission for sound encoding and predicts elevated auditory thresholds which we tested by recordings of auditory brainstem responses.

**Figure 6.**
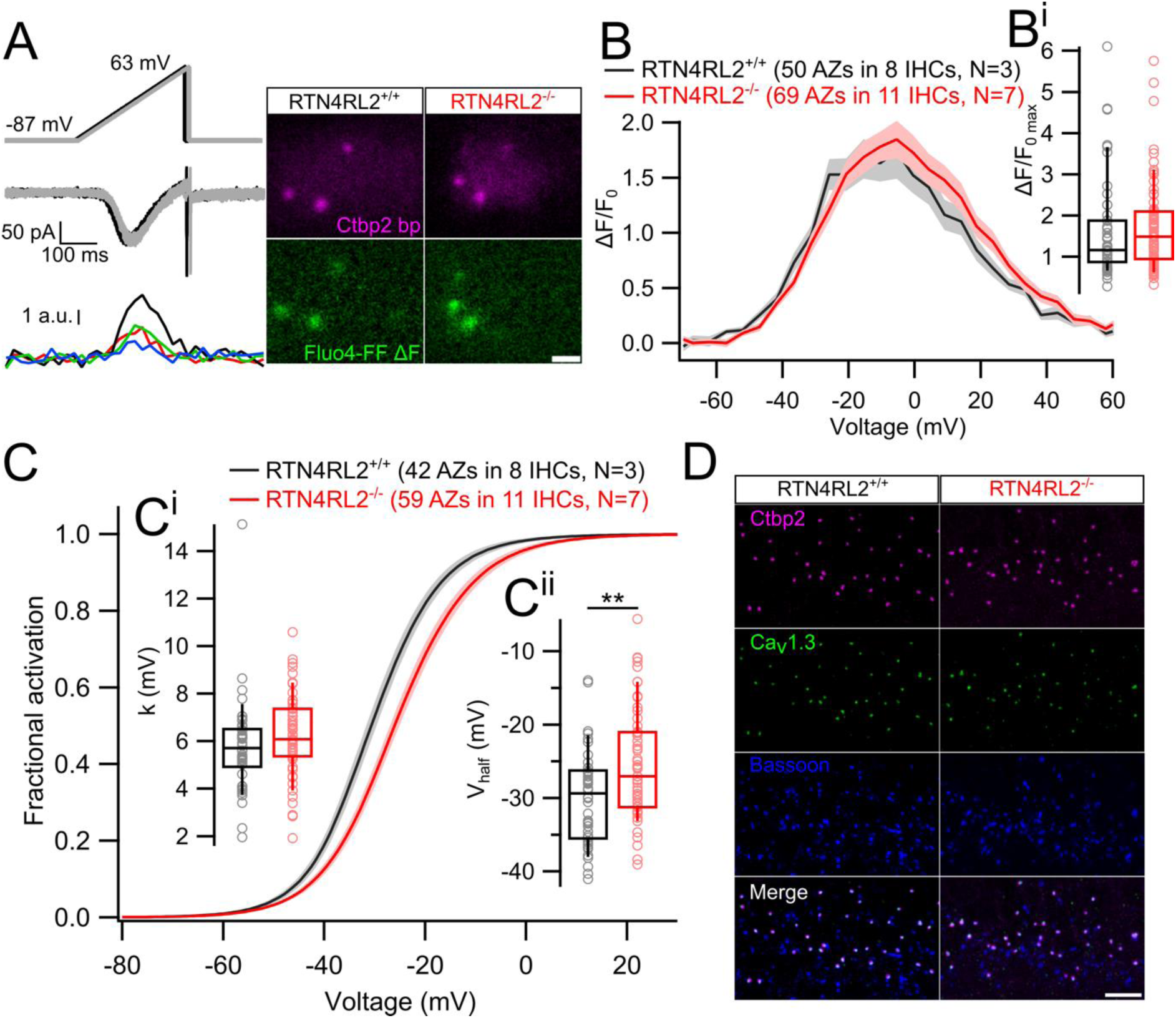
Depolarized shift of Ca^2+^ channel activation at single AZs but intact presynaptic organization in RTN4RL2^-/-^ IHCs. (**A**) Voltage ramp stimulation protocols (top), evoked whole cell currents (middle) and the presynaptic hotspots of Fluo4-FF fluorescence (bottom) of a representative IHC recording. Black and grey colors represent the two stimulations, one being 5 ms shifted over the other. Images on the right show single imaging planes of representative RTN4RL2^+/+^ (left) and RTN4RL2^-/-^ (right) IHCs filled with TAMRA conjugated Ctbp2 binding peptide (Ctbp2 bp) and Fluo4-FF Ca^2+^ dye. Ca^2+^ hotspots are visualized by subtracting the average of baseline planes from the average of 5 planes during stimulation. Scale bar = 2 μm. (**B**) Average fluorescence-voltage relationships of Ca^2+^ influx at single AZ from RTN4RL2^+/+^ and RTN4RL2^-/-^ IHCs show no difference in the maximal Ca^2+^ amplitude (**B^i^**; RTN4RL2^+/+^: 1.6 ± 0.17, SD = 1.2, n = 50 AZs vs RTN4RL2^-/-^: 1.7 ± 0.13 pA, SD = 1.09, n = 69 AZs; p = 0.24, Mann-Whitney-Wilcoxon test). Shaded areas represent ± SEM. (**C**) Average fractional activation curves of Ca^2+^ channels at single AZs show intact voltage sensitivity (**C^i^**; RTN4RL2^+/+^: 5.79 ± 0.32 mV, SD = 2.04 mV, n = 42 AZs vs RTN4RL2^-/-^: 6.25 ± 0.22 mV, SD = 1.7 mV, n = 59 AZs; p = 0.06, Mann-Whitney-Wilcoxon test) but depolarized shift of V_half_ (**C^ii^**; RTN4RL2^+/+^: -30 ± 1 mV, SD = 6.5 mV, n = 42 AZs vs RTN4RL2^-/-^: -25.5 ± 0.98 mV, SD = 7.49 mV, n = 59 AZs; p = 0.002, Student’s t-test) in RTN4RL2^-/-^ IHCs. Shaded areas represent ± SEM. (**D**) Representative immunolabelings of presynaptic proteins show no apparent mislocalization in RTN4RL2^-/-^ IHCs. Scale bar = 5 μm. Box-whisker plots show the median, 25/75 percentiles (box) and 10/90 percentiles (whiskers). Individual data points are overlaid.

### RTN4RL2 is essential for normal hearing

In order to evaluate auditory systems function in RTN4RL2^-/-^ mice, we recorded auditory brainstem responses (ABRs) at the age of 2-4 months. ABRs were elicited in response to 4, 8, 16, 32 kHz tone bursts and click stimuli of increasing sound pressure levels. We determined the sound thresholds by identifying the lowest sound pressure level which resulted in detectable ABR waveform. We found ABR thresholds to be significantly increased by around 30-45 dB across all the tested frequencies and in response to the click stimulations in RTN4RL2^-/-^ mice compared to the wild-type controls (Figure 7). RTN4RL2^+/-^ mice showed an intermediate phenotype between the RTN4RL2^-/-^ and RTN4RL2^+/+^ mice with a lower yet significant increase of ABR thresholds of approximately 10-15 dB at 4, 16 and 32 kHz frequencies.

**Figure 7.**
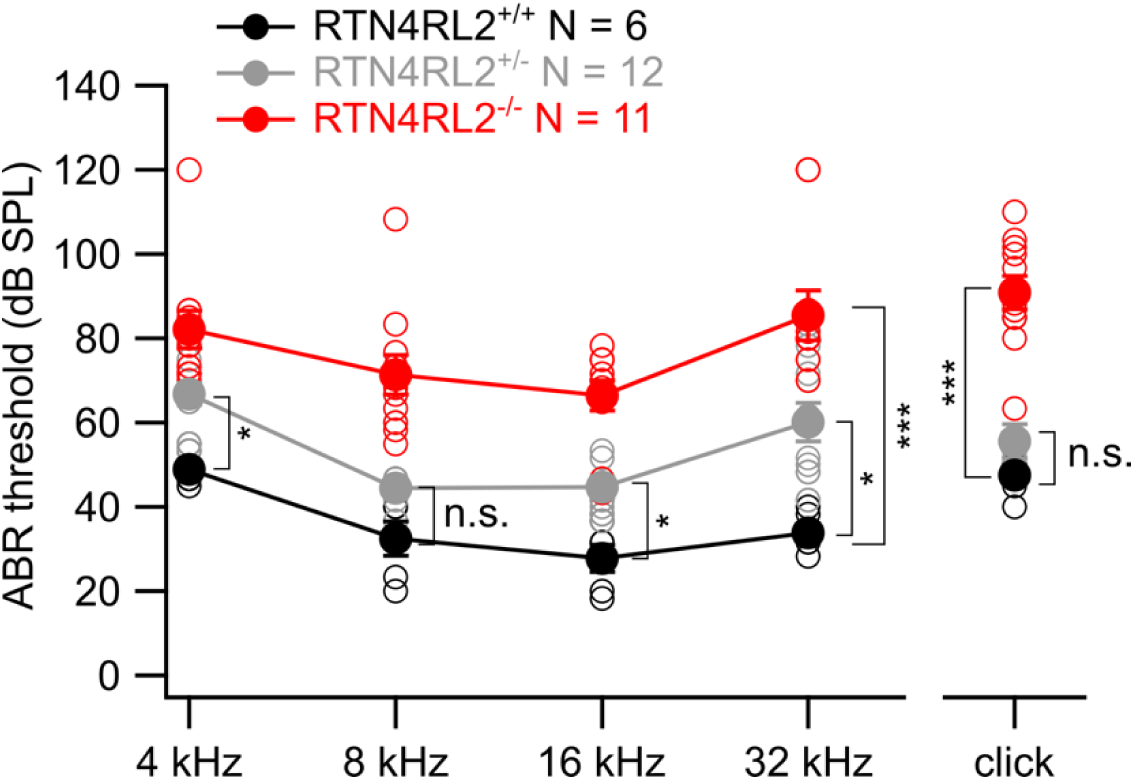
Elevated acoustic thresholds in RTN4RL2^-/-^ mice. ABR thresholds were measured in response to 4, 8, 16, 32 kHz tone bursts and click stimuli. ABR thresholds of individual animals are shown in open circles on top of the mean ± SEM. Statistical significances are reported as *p < 0.05, ***p < 0.001, Kruskal-Wallis followed by Dunn’s multiple comparison test.

## Discussion

In this study, we investigated the role of the Nogo/RTN4 receptor homolog RTN4RL2 in the cochlea. Consistent with the previous reports, we detected RTN4RL2 expression in IHCs and SGNs. Upon RTN4RL2 deletion, we observed alterations of the IHC-SGN synapses (Figure 8). Presynaptically, Ca^2+^ channel activation was shifted towards depolarized voltages and ribbon size was enlarged. IHC exocytosis was unaltered when probed at saturating depolarizations. At the postsynaptic side, PSDs juxtaposing presynaptic ribbons were smaller compared to the controls and seemed to be deficient of GluA2-4, despite the expression of Gria2 mRNA in RTN4RL2^-/-^ SGNs. Additionally, we observed PSDs which resided further from the IHC membrane and did not juxtapose the presynaptic AZs in RTN4RL2^-/-^ mice. These PSDs potentially belong to the type I SGN neurites that ended in the inner spiral bundle without contacting the IHCs, as observed in SBEM reconstructions of RTN4RL2^-/-^ SGNs. Finally, ABR thresholds were elevated in RTN4RL2^-/-^ mice indicating that RTN4RL2 is required for normal hearing.

**Figure 8.**
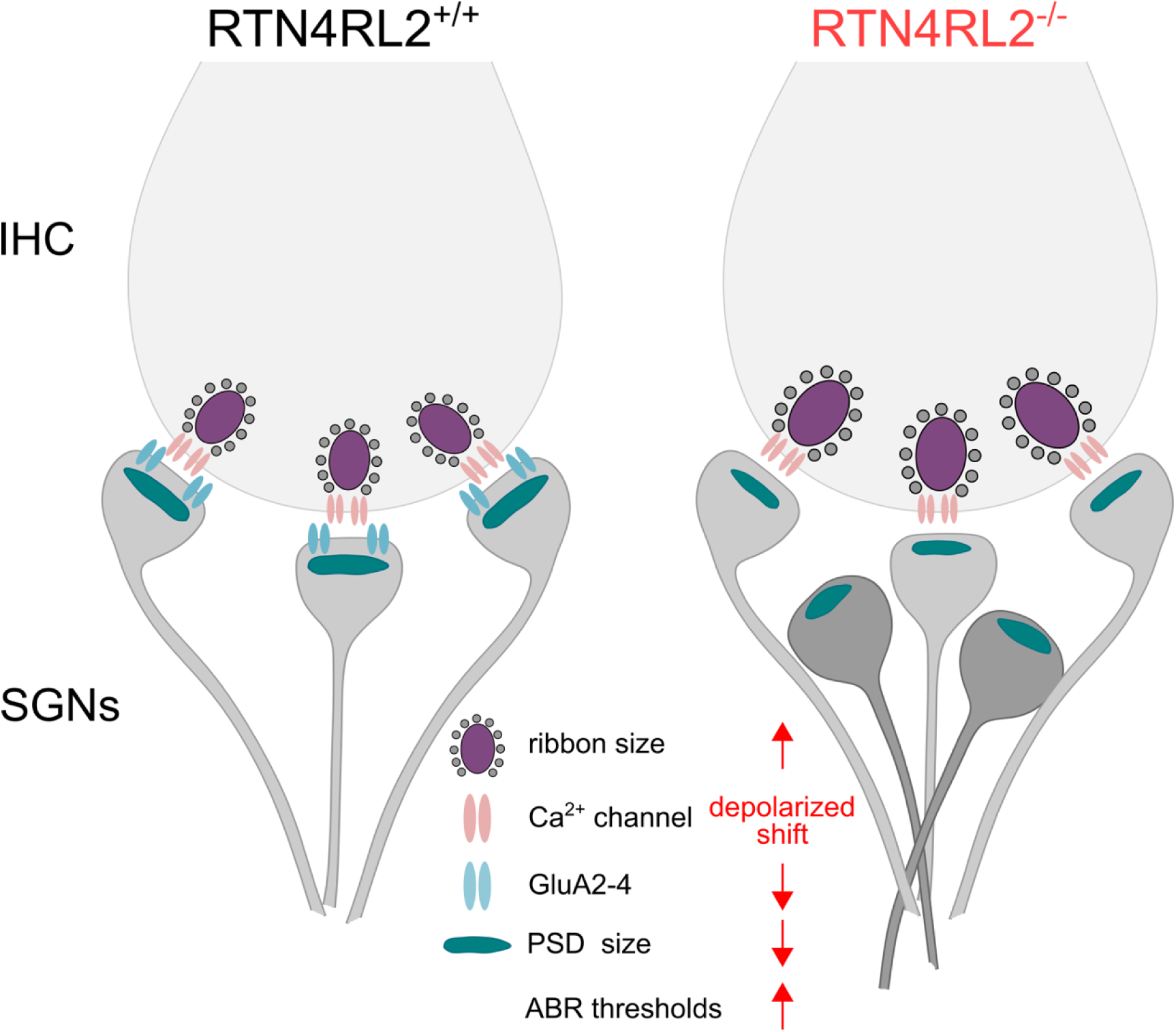
Schematic illustration of the key structural and functional changes in the auditory periphery of RTN4RL2^-/-^ mice. RTN4RL2^-/-^ mice display enlarged synaptic ribbons and depolarized shift in the activation of presynaptic Ca^2+^ channels in IHCs, as well as reduced size of PSDs juxtaposing presynaptic ribbons. RTN4RL2 deficiency further leads to a decrease in GluA2-4 AMPA receptor subunits at PSDs and results in a subset of type I SGN neurites that reach the inner spiral bundle but do not engage the IHCs.

RTN4 receptors have been implicated as presynaptic adhesion molecules based on their transsynaptic interactions with postsynaptically enriched brain-specific angiogenesis inhibitor (BAI) adhesion GPCRs (Stephenson et al., 2013; Wang et al., 2021). Despite this, at least RTN4R has been proposed to function postsynaptically, given that its knockdown leads primarily to postsynaptic changes (Wills et al., 2012). The interpretation of RTN4R function and localization is further complicated by their enrichment in both pre- and postsynaptic terminals (Wang et al., 2002b). Similarly, we observed RTN4RL2 expression in both IHCs and SGNs. Interestingly, a recent study described a hearing impairment in BAI1 deficient mice and suggested that postsynaptically functioning BAI1 is important for correct localization of AMPA receptor subunits in spiral ganglion neurons based on the absence or decrease of different AMPA receptor subunits at the SGN postsynapses in BAI1 deficient mice (Carlton et al., 2024). Similarly, our data from RTN4RL2^-/-^ mice show the absence of GluA2-4 positive puncta juxtaposing presynaptic AZs. Importantly, the BAI1 knock-out model in the study by Carlton et al. eliminated specifically long isoform of BAI1, which contains extracellular thrombospondin repeats (TSRs) important for mediating transsynaptic interactions, while leaving she short isoform - containing the transmembrane repeats and intracellular domain of the receptor - intact (Wang et al., 2021; Carlton et al., 2024). Along with the evidence for transsynaptic interaction between RTN4Rs and BAIs (Wang et al., 2021), this raises the possibility that the GluA2-4 AMPA receptor subunit localization to SGN PSDs requires the interaction of presynaptic RTN4RL2 and postsynaptic BAI1. Likewise, the changes of presynaptic Ca^2+^ channel properties and ribbons in IHCs could result from direct presynaptic action of RTN4RL2. Alternatively, RTN4RL2 could act postsynaptically in SGNs with an impact on transsynaptic signaling to IHCs. Presynaptic changes upon postsynaptic manipulation have been reported previously. For instance, the deletion of postsynaptically expressed Pou4f1 transcription factor resulted in hyperpolarized shift of IHC Ca^2+^ channels, as well as in changes of the presynaptic heterogeneity of IHC AZs (Sherrill et al., 2019). Similarly, presynaptic ribbon sizes were altered in IHCs of mice, which exhibited conditional deletion of Runx1 transcription factor from SGNs (Shrestha et al., 2023). Further experiments, employing mouse mutants with specific RTN4RL2 deletion in IHCs or SGNs will allow to evaluate the precise presynaptic and postsynaptic roles of RTN4RL2 including a possible transsynaptic signaling to IHC AZs.

We propose that the afferent synaptic changes contribute to the severe hearing impairment in RTN4RL2^-/-^ mice. The phenotype of increased ABR thresholds could partially be explained by the depolarized activation of Ca^2+^ channels in IHCs. As it has been shown, the apparent Ca^2+^ dependence of glutamate release from IHCs on average follows a near linear relationship (Wong et al., 2014; Özçete and Moser, 2021; Jaime Tobón and Moser, 2023b, 2023a). Therefore, due to the depolarized shift of the Ca^2+^ channel activation, the glutamate release from IHCs would require stronger stimuli. While exocytosis, probed by depolarizing the IHCs to -17 mV and recording cell membrane capacitance changes, remained intact in RTN4RL2^-/-^ mice, it could still be affected at milder depolarizations (e.g., -50 mV to -40 mV), which correspond to the receptor potential range of the IHCs. This could result in increased sound thresholds, as shown in Ribeye knock-out mice, where depolarized activation of Ca^2+^ channels has been associated with higher firing thresholds and lower spontaneous rates in SGNs, although a less severe hearing phenotype was observed in those mice (Jean et al., 2018). Furthermore, we have shown recently that a direct manipulation of the voltage dependence of Ca_v_1.3 Ca^2+^ channel activation results in changes of SGN firing and potentially auditory thresholds, further supporting our hypothesis of sound threshold elevation in RTN4RL2^-/-^ mice being caused by the depolarized activation of Ca^2+^ channels in IHCs (Karagulyan et al., 2025).

In addition or alternatively, the increase in auditory thresholds could be caused by the deficiency of GluA2-4 AMPA receptors of the SGN postsynapses in RTN4RL2^-/-^ mice. Compared to GluA3 KO mice, which display reduced GluA2 signal at SGN PSDs but intact auditory thresholds, the increased auditory thresholds in RTN4RL2^-/-^ mice could additionally be caused by the lack of GluA4 subunits juxtaposing presynaptic ribbons, which, in contrast, were shown to be upregulated in GluA3 KO mice (Rutherford et al., 2023). Given the observed synaptic changes, it seems less likely that the hearing impairment would have resulted from the intrinsic changes of SGN firing. Further experiments, where SGNs would be stimulated directly via *ex vivo* patch-clamp or optogenetically could help to probe for functional deficits in SGNs of RTN4RL2^-/-^ mice. Another interesting observation was the additional PSDs not juxtaposing presynaptic ribbons that were detected around IHCs of RTN4RL2^-/-^ mice. These “orphan” PSDs seem not to be part of the efferent synapses formed between LOC/MOCs and SGNs as we verified by simultaneous anti-Homer1 and anti-synapsin immunolabelings. To check the SGN connectivity we performed SBEM in RTN4RL2^+/+^ and RTN4RL2^-/-^ cochlear samples and observed a high percentage of non-synaptic SGN neurites in the reconstructed neurons of inner spiral bundles in RTN4RL2^-/-^ samples. A sizable subset of the non-synaptic SGN boutons did not make a contact with IHCs, potentially housing the “orphan” PSDs observed around the IHCs in our immunolabeled samples. How exactly those non-synaptic neurites appear remains to be investigated. SGN neurite retraction from IHCs has been demonstrated in noise-exposed animals, followed by the degeneration of the SGN somata and axons (Kujawa and Liberman, 2009, 2015; Moverman et al., 2023). This process however results in up to 50% degeneration of the ribbon synapses. Interestingly, in RTN4RL2^-/-^ mice the number of the ribbon synapses as well as the density of the SGN somata was largely intact. This indicates that the non-synaptic neurites do not result from synaptic loss as found in hereditary or acquired synaptopathy (Roux et al., 2006; Ruel et al., 2008; Seal et al., 2008; Kujawa and Liberman, 2009). Although it is plausible that the non-synaptic SGN neurites may fail to reach the IHCs during initial development, it is unlikely that this would selectively impact only a subset of neurons and still allow the Homer1-positive patches to develop in the absence of presynaptic components. Alternatively, it is known that major synaptic pruning takes place in developing SGNs, whereby up to 50% of the synapses are lost between the IHCs and the SGNs (Defourny et al., 2013; Coate et al., 2019), suggesting that the non-synaptic SGN neurites of RTN4RL2^-/-^ mice could be a result of failed pruning of SGNs during the development. Yet how this would then comply with normal synapse and SGN counts remains to be elucidated. Future studies will be required to understand the fate of type I SGNs in RTN4RL2^-/-^ mice during development by mapping their full-length periphery projections with large scale imaging tools.

Future work will be required to identify the RTN4RL2 ligands in the cochlea. To this date, the interaction partners of RTN4Rs are not very well understood. While RTN4R in-trans interactions include NogoA, MAG, OMgp, chondroitin sulfate proteoglycans, the only well established ligand for RTN4RL2 remains MAG (Fournier et al., 2001; Domeniconi et al., 2002; Wang et al., 2002a; Venkatesh et al., 2005; Schwab, 2010). Other attractive candidate ligands for RTN4Rs are the BAIs. BAI-RTN4R interaction has been suggested to mediate dendritic arborization, axonal elongation, synapse formation in IPSCs (Wang et al., 2021) and, in the case of this study GluA2-4 AMPA receptor subunit localization to the SGN postsynaptic densities. Finally, RTN4RL2 has been proposed to interact with chondroitin-sulfate proteoglycan Versican to control the amount of skin innervation by DRG neurites (Bäumer et al., 2014). While to our knowledge, attempts have yet to be made for detecting Versican signal around the IHCs, other extracellular matrix proteins such as aggrecan, brevican, Tenascin-R, Tenascin-C, hyaluron and proteoglycan link proteins 1 and 4 have been shown to form dense, basket-like structures that surround the base of the IHCs (Kwiatkowska et al., 2016; Sonntag et al., 2018). Furthermore, deletion of brevican results in alteration of pre- and postsynaptic spatial coupling at IHC synapses, indicating the possible role of extracellular matrix in transsynaptic organization. However, while the existing observations suggest that the synaptic changes in RTN4RL2 deficient animals might reflect derailed interaction of the SGN neurites with the extracellular matrix, the relevance of such an interaction of RTN4RL2 and Versican in the cochlea needs to be addressed in the future.

## Materials and Methods

### Animals

RTN4RL2 null mutant mice on a C57BL/6N background were generated as previously described (Wörter et al., 2009). Knock-out (RTN4RL2^-/-^) and wild-type (RTN4RL2^+/+^) mice were derived from heterozygous matings. Both male and female mice were used in this study. Animals were genotyped as previously described (Wörter et al., 2009). The ages of the mice varied depending on the experiment as noted in the manuscript. The breeding and the experiments were approved by the Institutional Animal Care and Use Committee at the Medical University of Innsbruck, local Animal Welfare Committee of the University Medical Center Göttingen and the Max Planck Institute for Multidisciplinary Sciences, as well as the Animal Welfare Office of the state of Lower Saxony, Germany (LAVES, AZ: 19/3134). The ABR experiments were carried out under the approval of the Austrian Ministry of Education, Science and Research (reference number BMWFW-66.011/0120-WF/V/3b/2016).

### Patch-clamp recordings

The apical turn of the organ of Corti was dissected from p21 to p29 animals in HEPES Hanks solution containing (in mM): 5.36 KCl, 141.7 NaCl, 10 HEPES, 0.5 MgSO_4_, 1 MgCl_2_, 5.6 D-glucose, and 3.4 L-glutamine (pH 7.2, ∼ 300 mOsm/l). IHCs were exposed by gently removing of nearby supporting cells by negative pressure through a glass pipette from either modiolar or pillar side. All experiments were conducted at room temperature (RT, 20–25°C). Patch pipettes were made from GB150-8P or GB150F-8P borosilicate glass capillaries (Science Products, Hofheim, Germany) for perforated and ruptured patch-clamp configurations, respectively. Pipettes were fire polished with a custom-made microforge and coated with Sylgard to minimize the capacitive noise, whenever capacitance recordings were performed. The measurements were performed using EPC-10 amplifier controlled by Patchmaster software (HEKA Elektronik, Germany). The holding potential for IHCs was set to -87 mV across all the experiments.

### Ruptured patch-clamp

For ruptured patch-clamp we used extracellular solution containing (in mM): 2.8 KCl, 105 NaCl, 10 HEPES, 1 CsCl, 1 MgCl_2_, 5 CaCl_2_, 35 TEA-Cl, and 2 mg/ml D-glucose (pH 7.2, ∼ 300 mOsm/l).

Intracellular solution contained in mM: 111 Cs-glutamate, 1 MgCl_2_, 1 CaCl_2_, 10 EGTA, 13 TEA-Cl, 20 HEPES, 4 Mg-ATP, 0.3 Na-GTP and 1 L-glutathione (pH 7.3, ∼ 290 mOsm/l). Additionally, Ca^2+^ indicator Fluo4-FF (0.8 mM, Life Technologies) and TAMRA-conjugated RIBEYE/Ctbp2 binding peptide (10 mM, synthetized by the group of Dr. Olaf Jahn, Göttingen) were added to the intracellular solution for live imaging (Zenisek et al., 2004). Voltage dependency of Ca^2+^ influx was recorded by applying 20 ms long step deploarizations with 5 mV increment. For Ca^2+^ imaging, voltage ramp depolarizations ranging from -87 mV to 63 mV in the course of 150 ms were applied to the cells. Leak correction was performed using p/4 protocol and liquid junction potential of 17 mV was corrected offline. Recordings were discarded from the analysis if the series resistance (R_s_) exceeded 14 MOhm during the first 3 minutes after rupturing the cell, leak current exceeded -50 pA at the holding potential and Ca^2+^ current rundown was more than 25%.

Recordings were analyzed using Igor Pro 6.3 (Wavemetrics) custom-written programs. Ca^2+^ current-voltage relationships (IV curves) were obtained by averaging approximately 5 ms segments at the maximal activation regions of individual current traces and plotting them against the depolarization voltages. Fractional activation curves of the Ca^2+^ channels were calculated from the IVs, normalized and fitted with the Boltzmann function: 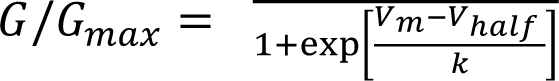, where *G* is conductance, *G_max_* is the maximal conductance, *V_m_* is the membrane potential, *V_half_* is the voltage corresponding to the half maximal activation of Ca^2+^ channels and *k* (slope of the curve) is the voltage sensitivity of Ca^2+^ channel activation.

### Perforated patch clamp recordings

To perform simultaneous membrane capacitance (C_m_) and Ca^2+^ current measurements we used extracellular solution containing (in mM): 106 NaCl, 35 TEA-Cl, 2.8 KCl, 1 MgCl_2_, 1 CsCl, 10 HEPES, 3 CaCl_2_, and 5.6 D-glucose (pH 7.2, ∼ 300 mOsm/l). The pipette solution contained (in mM): 137 Cs-gluconate, 15 TEA-Cl, 3 GTP-Na_2_, 1 ATP-Mg, 10 HEPES, 1 MgCl_2_, as well as 300 mg/ml amphotericin B (pH 7.17, ∼ 290 mOsm/l). Perforated patch-clamp recordings from IHCs were described previously (Moser and Beutner, 2000). C_m_ changes were measured using Lindau-Neher technique (Lindau and Neher, 1988). To evoke exocytosis IHCs were stimulated by step depolarizations (to -17 mV) of different durations (2, 5, 10, 20, 50, 100 ms) in randomized manner. Currents were leak-corrected using a p/5 protocol. Recordings were used only if the leak current was lower than 30 pA and the R_s_ was lower than 30 MOhm. Recordings were analyzed using Igor Pro 6.3 (Wavemetrics) custom-written programs. Ca^2+^ charge (Q_Ca_^2+^) was calculated by the time integral of the leak-subtracted Ca^2+^ current during the depolarization step. ΔC_m_ was calculated as the difference between the average C_m_ before and after the depolarization. To measure average C_m_ we used 400 ms segments and skipped the initial 100 ms after the depolarization.

### Ca^2+^ imaging

Ca^2+^ imaging was performed using spinning disc confocal microscope, as described before (Ohn et al., 2016). Briefly, the set-up was equipped with a spinning disc confocal unit (CSU22, Yokogawa) mounted on an upright microscope (Axio Examiner, Zeiss) and scientific CMOS camera (Andor Neo). Images were acquired using 63x, 1.0 NA objective (W Plan-Apochromat, Zeiss). The pixel size was measured to be 103 nm.

IHCs were loaded with Fluo4-FF Ca^2+^ dye and TAMRA-conjugated Ctbp2 binding dimeric peptide via the patch pipette. First, the cells were scanned from bottom to top by imaging TAMRA fluorescence with 561 nm laser (Jive, Cobolt AB) and exposing each plane for 0.5 seconds. The stack was acquired with 0.5 μm step size using Piezo positioner (Piezosystem). This allowed us to identify the planes containing synaptic ribbons. Next, we recorded Fluo4- FF fluorescence increase at individual synapses by imaging ribbon containing planes with 491 nm laser (Calypso, Cobolt AB) at 100 Hz while applying voltage ramp depolarizations to the cell. 2 voltage ramps were applied at each plane, one being 5 ms shifted relative to the other. Spinning disk rotation speed was set to 2000 rpm, in order to synchronize with Ca^2+^ imaging. Images were analyzed using Igor Pro 6.3 (Wavemetrics). Ca^2+^ hotspots were identified by subtracting the average signal of several baseline frames from the average signal of 5 frames during stimulation (ΔF image). Since the same Ca^2+^ hotspot appears across multiple planes, the plane exhibiting the strongest signal was chosen. The intensities of the 3x3 matrix surrounding the central pixel of the hotspot were averaged across all time points to obtain the intensity profiles of Ca^2+^ influx over time. Afterwards, the background signal (average of approximately 60x60 pixel intensities outside the cell) was subtracted from the intensity-time profiles and ΔF/F_0_ traces were calculated. To enhance voltage resolution of Ca^2+^ imaging, two ΔF/F_0_ traces corresponding to two voltage ramp depolarizations (one shifted by 5 ms over the other) were combined and plotted against the corresponding voltages (FV curves). These curves were subsequently fitted with a modified Boltzmann function. Afterwards, fractional activation curves were calculated by fitting the linear decay of the fluorescence signal from the FV curves with a linear function (G_max_) and then dividing the FV fit by the G_max_ line. The resulting curves were further fitted with a Boltzmann function. Maximal Ca^2+^ influx (ΔF/F_0 max_) was calculated by averaging 5 points during the stimulation.

### Immunohistochemistry

For whole mount immunofluorescence, cochleae were fixed in 4% formaldehyde on ice for 45-60 minutes or sometimes overnight. For anti-Ca_V_1.3 stainings the cochleae were fixed for 10 minutes. After the fixation, the organs of Corti were microdissected in PBS and blocked in GSDB (goat serum dilution buffer; 16% goat serum, 20 mM phosphate buffer (PB), 0.3% Triton X-100, 0.45 M NaCl) for 1 hour at RT. The samples were then incubated in the primary antibody mixture overnight at 4°C. For anti-GluA4 immunolabelings samples were incubated at primary antibody mixture at 37°C overnight. The next day, samples were washed 3 times using wash buffer (20 mM PB, 0.3% Triton X-100, 0.45 M NaCl) followed by incubation in the secondary antibody mixture for 1 hour at RT. Afterwards, the samples were washed 3 times in the washing buffer, one time in PB and mounted using mounting medium (Mowiol).

The following primary antibodies were used (Table 1): rabbit anti-Homer1 (1:500, 160 002, Synaptic Systems), mouse anti-Ctbp2 (1:200, 612044, BD Biosciences), rabbit anti-Ca_V_1.3 (1:100, ACC-005, Alomone Labs), mouse anti-Bassoon (1:300, ab82958, Abcam), mouse anti GluA2 (1:200, MAB397, Millipore), rabbit anti GluA4 (1:200, AB1508, Millipore), rabbit anti GluA2/3 (1:200, AB1506, Chemicon), rabbit anti-Myosin7a (1:800, ab3481, Abcam, or Proteus BioSciences), guinea pig anti-RibeyeA (1:500, 192104, Synaptic Systems), guinea pig anti- Synapsin1/2 (1:500, 106004, Synaptic Systems), guinea pig anti-Vglut3 (1:500, 135204, Synaptic Systems), chicken anti-parvalbumin (1:200, 195006, Synaptic Systems), chicken anti-calretinin (1:200, 214106, Synaptic Systems), guinea pig anti-VAChT (1:1000, 139105, Synaptic Systems), mouse anti-ATP1A3 (1:300, MA3-915, Thermo Fisher Scientific).

**Table 1.**
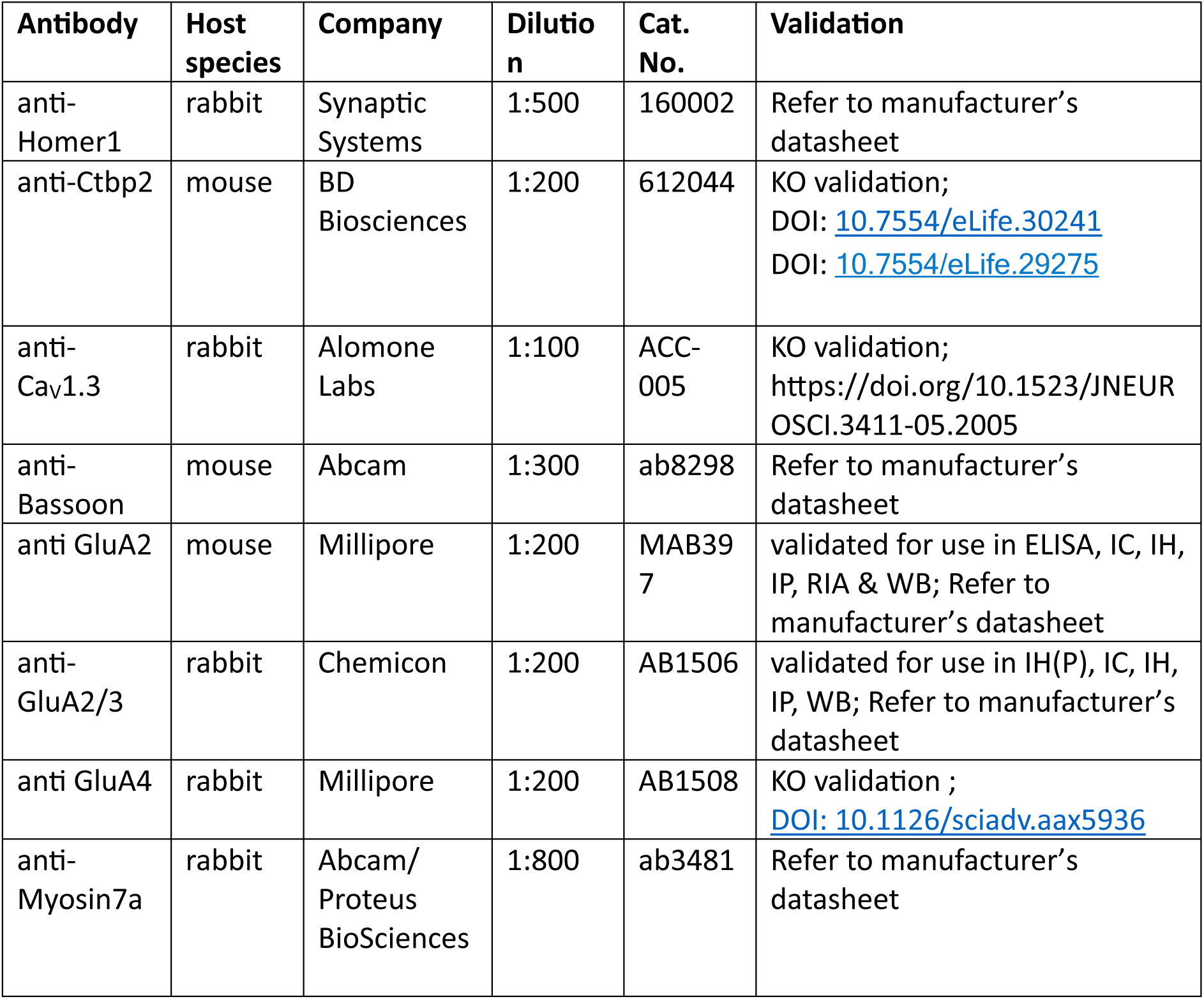
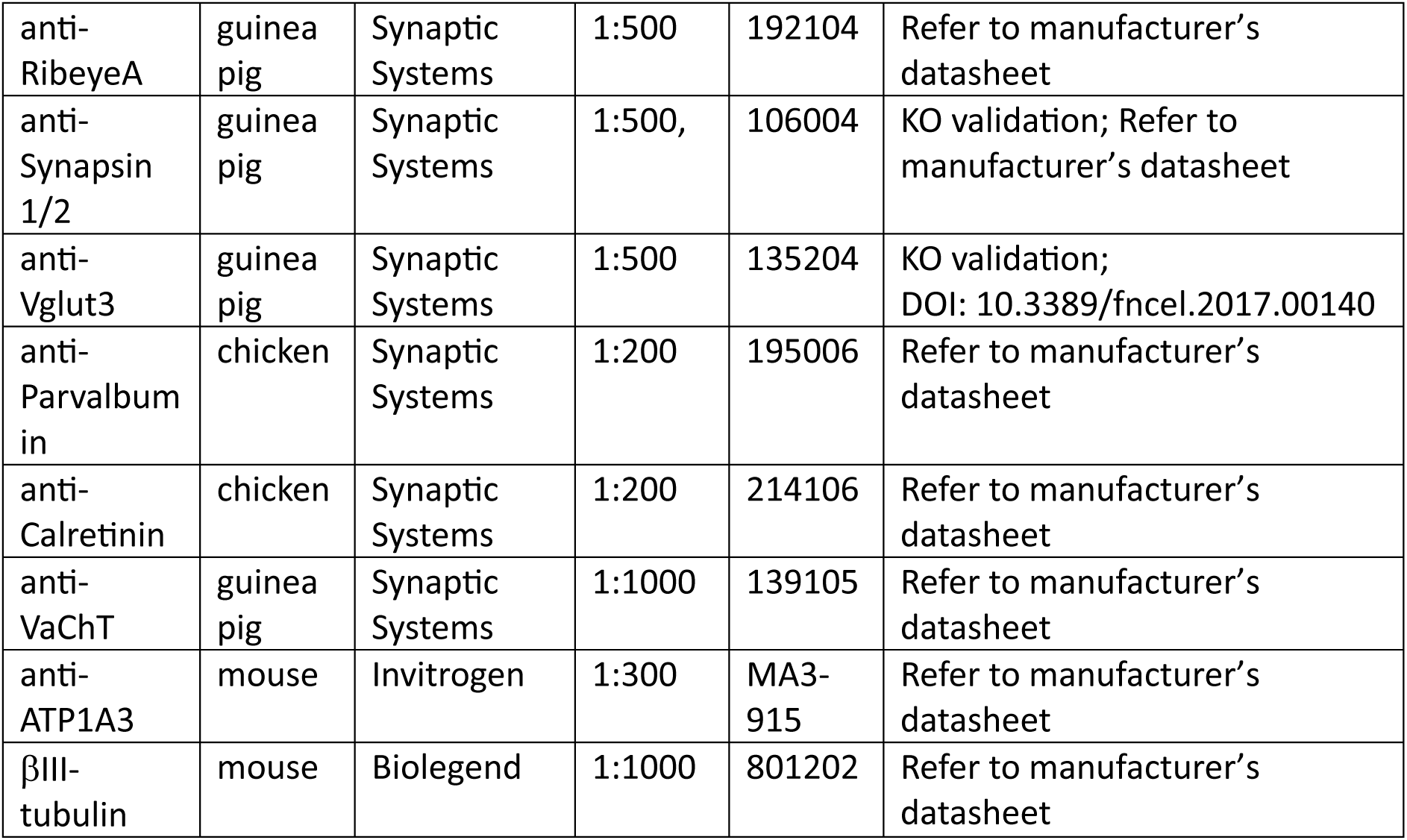
List of primary antibodies.

The following secondary antibodies were used: Alexa Fluor 488 conjugated anti-guinea pig (1:200, A11073, Thermo Fisher Scientific), Alexa Fluor 488 conjugated anti-rabbit (1:200, A11008, Thermo Fisher Scientific), Alexa Fluor 568 conjugated anti-chicken (1:200, ab175711, Abcam), Alexa Fluor 633 conjugated anti-guinea pig (1:200, A21105, Thermo Fisher Scientific), STAR 580 conjugated anti-mouse (1:200, Abberior, ST635P-1001-500UG), STAR 635 conjugated anti-rabbit (1:200, Abberior, ST635P-1002-500UG).

Confocal stacks were acquired using Leica SP8 confocal microscope or Abberior Instruments Expert Line STED microscope.

The volumes of the Ctbp2 and Homer1 positive puncta were estimated using the surface algorithm of Imaris software (version 9.6.0, Bitplane). Following parameters were used to create the surfaces for both Ctbp2 and Homer1 puncta in all the analyzed stacks: surface detail 0.07 μm, background subtraction 0.562 μm, touching object size 0.4 μm.

The brightness and the contrast of the representative images were adjusted using Fiji (ImageJ) software for the visualization purposes.

### Spiral ganglion neuron densities

RTN4RL2^+/+^ and RTN4RL2^-/-^ cochlea from p40 mice of either sex were processed for SGN counts. Mouse monoclonal anti-βIII-tubulin (1:1000, 801202, Biolegend) at 4°C, followed by secondary Alexa Fluor 488 conjugated anti-mouse (1:1000, Thermo Fisher Scientific), was used to label all SGNs. Nuclei were visualized using a DAPI fluorescent counterstain (1:1000, Life Technologies). 20X images of mid-modiolar cochlear sections were collected. ImageJ was used to outline the spiral ganglion and generate the area. Quantitative assessment was performed on every other mid-modiolar section to reduce chances of double counting. Labeled cells were counted only if they had a round cell body, presence of nucleus, and homogenous cytoplasm. Densities were calculated and statistical differences were measure using a Student’s t-test (GraphPad Prism, GraphPad Software Inc.).

### Inner and outer Hair cell counts

IHC and OHC counts were performed for RTN4RL2^+/+^ and RTN4RL2^-/-^ mice at the ages of p15, 1 month and 2 months. Images were imported to ImageJ software, where IHC/OHC counts were performed blinded to genotype using the ‘Cell counter’ tool. To quantify the cells the nuclear staining of the DAPI channel was used. The number of IHCs and OHCs was assessed along a length of 100 μm for each image, and then averaged across all samples to obtain the average number of IHCs (or OHCs) per 100 μm.

### RNAscope

RNA *in-situ* hybridization was performed using RNAscope^®^ (Advanced Cell Diagnostics). Standard mouse probes were used to examine the expression of RTN4RL2 (450761, ACD) and Gria2 (865091, ACD) closely following the manufacturer’s instructions. Briefly, 6 mm paraffin tissue sections underwent deparaffinization with xylene and a series of ethanol washes. Tissues were heated in kit-provided antigen retrieval buffer and digested by kit-provided proteinase. Sections were exposed to mFISH target probes and incubated at 40°C in a hybridization oven for 2 hours. After rinsing, mFISH signal was amplified using company-provided pre-amplifier and amplifier conjugated to fluorescent dye. Subsequently, sections were blocked with 1% BSA, 2% normal goat serum in 1xPBS containing 0.3% Triton X-100 for 1 hour at RT. The tissue was incubated in mouse anti-βIII-tubulin antibody (1:500, 801202, Biolegend) overnight at 4°C. The next day, sections were rinsed with PBS, blocked again before incubating in secondary Alexa Fluor 488 conjugated anti-mouse antibody (1:1000, Invitrogen) for 2 hours at RT. Sections were counterstained with DAPI (1:1000, Life Technologies), mounted, and stored at 4°C until image analysis. mFISH images were captured on a Leica SP8 confocal microscope and processed using ImageJ.

### Serial block-face scanning electron microscopy (SBEM)

#### Sample preparation for SBEM

One RTN4RL2^+/+^ and two RTN4RL2^-/-^ female mice at p36 were used for the SBEM experiments. Cochleae were processed for SBEM imaging as previously described (Hua et al., 2021).

In brief, the animals were decapitated after CO_2_ inhalation under anesthesia. The cochleae were dissected from the skulls and immediately fixed by perfusing with ice-cold fixative through the round and oval windows at constant flow speed using an infusion pump (Micro4, WPI). The fixative solution was freshly prepared and made of 4% paraformaldehyde (Sigma-Aldrich), 2.5% glutaraldehyde (Sigma-Aldrich), and 0.08 M cacodylate (pH 7.4, Sigma-Aldrich). After being immersed in the fixative at 4°C for 5 hours, the cochleae were transferred to a decalcifying solution containing the same fixative and 5% ethylenediaminetetraacetic acid (EDTA, Serva) and incubated at 4°C for 5 hours. The samples were then washed twice with 0.15 M cacodylate (pH 7.4) for 30 min each, sequentially immersed in 2% OsO_4_ (Sigma-Aldrich), 2.5% ferrocyanide (Sigma-Aldrich), and again 2% OsO_4_ at RT for 2, 2, and 1.5 hours. After being washed in 0.15 M cacodylate and distilled water (Sartorius) for 30 min each, the samples were sequentially incubated in filtered 1% thiocarbohydrazide (TCH, Sigma-Aldrich) solution and 2% OsO_4_ at RT for 1 and 2 hours, as well as in lead aspartate solution (0.03 M, pH 5.0, adjusted with KOH) at 50 °C for 2 hours with immediate two washing steps with distilled water at RT for 30 min each. For embedding, the samples were dehydrated through graded pre-cooled acetone (Carl Roth) series (50%, 75%, 90%, for 30 min each, all cooled at 4 °C) and then pure acetone at RT (three times for 30 min each). The sample infiltration started with 1:1 and 1:2 mixtures of acetone and Spurr’s resin monomer (4.1 g ERL 4221, 0.95 g DER 736, 5.9 g NSA and 1% DMAE; Sigma-Aldrich) at RT for 6 and 12 hours on a rotator. After being impregnated in pure resin for 12 hours, the samples were placed in embedding molds (Polyscience) and hardened in a pre-warmed oven at 70°C for 72 hours.

#### Sample trimming and SBEM imaging

The sample blocks were mounted upright along the conical center axis on aluminum metal rivets (3VMRS12, Gatan, UK) and trimmed coronally towards the modiolus using a diamond trimmer (TRIM2, Leica, Germany). For each sample, a block face of about ∼ 600 x 800 mm^2^, centered at the apical segment based on the anatomical landmarks, was created using an ultramicrotome (UC7, Leica, Germany). The samples were coated with a 30 nm thick gold layer using a sputter coater (ACE600, Leica, Germany). The serial images were acquired using a field-emission scanning EM (Gemini300, Carl Zeiss, Germany) equipped with an in-chamber ultramicrotome (3ViewXP, Gatan, UK) and back-scattered electron detector (Onpoint, Gatan, UK). Focal charge compensation was set to 100 % with a high vacuum chamber pressure of 2.8 x 10^3^ mbar. The following parameters were set for the SBEM imaging: 12 nm pixel size, 50 nm nominal cutting thickness, 2 keV incident beam energy, and 1.5 ms pixel dwell time.

For the RTN4RL2^+/+^ dataset, 2377 consecutive slices (9000 × 15000 pixels) were collected, whereas the two RTN4RL2^-/-^ datasets had 3217 slices (16000 × 10000 pixels) and 2425 slices (9000 × 15000 pixels). All datasets were aligned along the z-direction using a custom MATLAB script based on cross-correlation maximum between consecutive slices (Hua et al., 2022) before being uploaded to webKnossos (Boergens et al., 2017) for skeleton and volume tracing.

#### Identification and quantification of auditory afferent fibers

In our SBEM datasets of the cochlea, manual skeleton tracing was carried out on all neurites that originated from three neighboring habenula perforata (HP) at the center of each dataset. To search for type I afferent fibers, several morphological features were used, such as myelination after entering HP, radial and unbranched fiber trajectory, contact with IHCs, as well as ribbon-associated terminals (Hua et al., 2021). This resulted in 115 putative type I afferent fibers and further classification was made based on the presence of presynaptic ribbon, fiber branching, and contact with IHC. For an illustration purpose, a type I afferent fiber bundle of one RTN4RL2^-/-^ dataset was volume traced and 3D rendered using Amira software (Thermo Scientific, US).

#### Ribbon size measurement and synapse counting

Ribbon-type synapses were manually annotated in 18 intact IHCs captured by SBEM using webKnossos. The dense core region of individual ribbon synapses was manually contoured, and the associated voxels were counted for ribbon volume measurement. In the case of multi-ribbon synapses, all ribbon bodies at a single active zone were summed up to yield the ribbon volume.

### Auditory brainstem responses

Auditory brainstem resposes (ABR) were recorded in 2- to 4-month-old mice of either sex, as described previously (Luque et al., 2021). Briefly, we used a custom-made system in an anechoic chamber in a calibrated open-field configuration. ABRs were recorded via needle electrodes in response to tone bursts of 4, 8, 16, and 32 kHz or a click wide spectrum (2-45,2 KHz, 2 KHz steps) as stimuli. Tone pips of 3 ms duration (1 ms rise and fall time) were presented at a rate of 60/s with alternating phases. Starting with 0 dB, the stimuli were increased in 5 dB steps up to 120 dB. Hearing thresholds were determined as the minimum stimulation level that produced a clearly recognizable potential. Both ears were tested. Since there was no significant difference between the right and left ears, we did not consider this factor further. Recordings were evaluated by three independent researchers in a blinded manner.

### Data analysis and statistics

The data were analyzed using Igor Pro (Wavemetrics), Python and GraphPad Prism (GraphPad Software Inc.) software. For 2 sample comparisons, data were tested for normality and equality of variances using Jarque-Bera and F-test, respectively. Afterwards, two-tailed Student’s t-test or Mann-Whitney-Wilcoxon test were performed. The latter was used when normality and/or equality of variances were not met. P-values were corrected for multiple comparisons using Holm-Šídák method. For ABR thresholds Kruskal-Wallis test followed by Dunn’s multiple comparison test was used. Data is presented as mean ± standard error of the mean (SEM), unless otherwise stated. Mean, SEM and standard deviation (SD) are indicated in the figure legends as mean ± SEM, SD. Significances are reported as *p < 0.05, **p < 0.01, ***p < 0.001. The number of the animals is indicated as N.

## Acknowledgements

We thank Sina Langer, Christiane Senger-Freitag, Sandra Gerke for expert technical support. We thank Prof. Dr. Carolin Wichmann, Dr. Susann Michanski, Julius Bahr and Sophia Mutschall for the help with the SBEM sample preparation. We thank Dr. Yi Jiang and Haoyu Wang for the visualization of neurite reconstruction. We further thank Dr. Mark Rutherford for providing tips for GluA4 immunolabeling. We would also like to thank Prof. Dr. Olaf Jahn and Lars van Werven for the TAMRA-conjugated Ctbp2 binding peptide synthesis. The study was supported by German Research Foundation through the Cluster of Excellence (EXC2067) Multiscale Bioimaging (TM) and the Leibniz Program (MO896/5 to TM) as well as by the European Research Council through the Advanced Grant “DynaHear” to TM under the European Union’s Horizon 2020 Research and Innovation program (grant agreement No. 101054467), and by Fondation Pour l’Audition (FPA RD-2020-10), by the SPIN-FWF grant to CB, and by the National Natural Science Foundation of China (82171133 to YH), Industrial Support Fund of Huangpu District .in Shanghai (XK2019011 to YH), Innovative Research Team of High-level Local Universities in Shanghai (SHSMU-ZLCX20211700). NK is a member of the Hertha Sponer College from the Cluster of Excellence Multiscale Bioimaging (MBExC). TM is a Max-Planck Fellow at the Max Planck Institute for Multidisciplinary Sciences. Open access funding provided by Max Planck Society.

## Author contributions

CB, TM, YH, and NK designed the research. NK, MÜ, YQ, NB, FH, LJC, FW, ML, RG performed experiments and analyzed data. NK and TM wrote the paper with contributions of YH, CB, YQ, NB. CB, TM, YH acquired funding. All authors have reviewed and approved the final version of the manuscript for submission.

## Supplementary Figures

**Figure 2-figure supplement 1.**
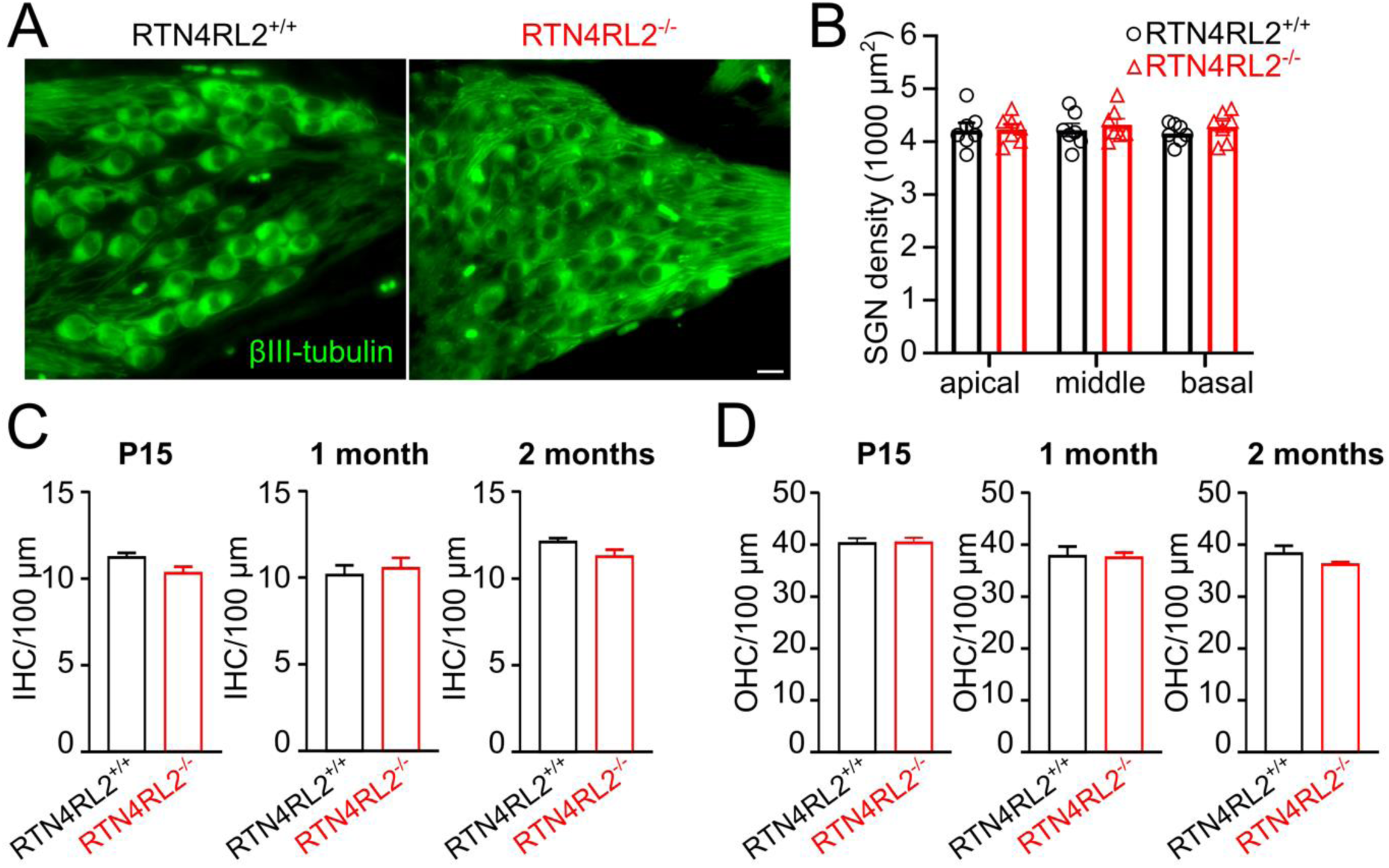
Cochlear cell densities are not changed in RTN4RL2^-/-^ mice. (**A**) Mid/Basal-modiolar sections labeled for βIII-tubulin (green, neurons) from p40 RTN4RL2^+/+^ and RTN4RL2^-/-^ mice. Exemplary section for RTN4RL2^-/-^ is the zoomed out image presented in figure 1C. (**B**) Quantitative analysis shows that the density of SGN cell bodies are similar between RTN4RL2^-/-^ and control cochleae. Data is presented as mean ± SD; N = 7 per group. Scale bar = 10 μm. (**C, D**) IHC (C) and OHC (D) densities are not affected in P15, 1 month and 2 months old RTN4RL2^-/-^ mice.

**Figure 2-figure supplement 2.**
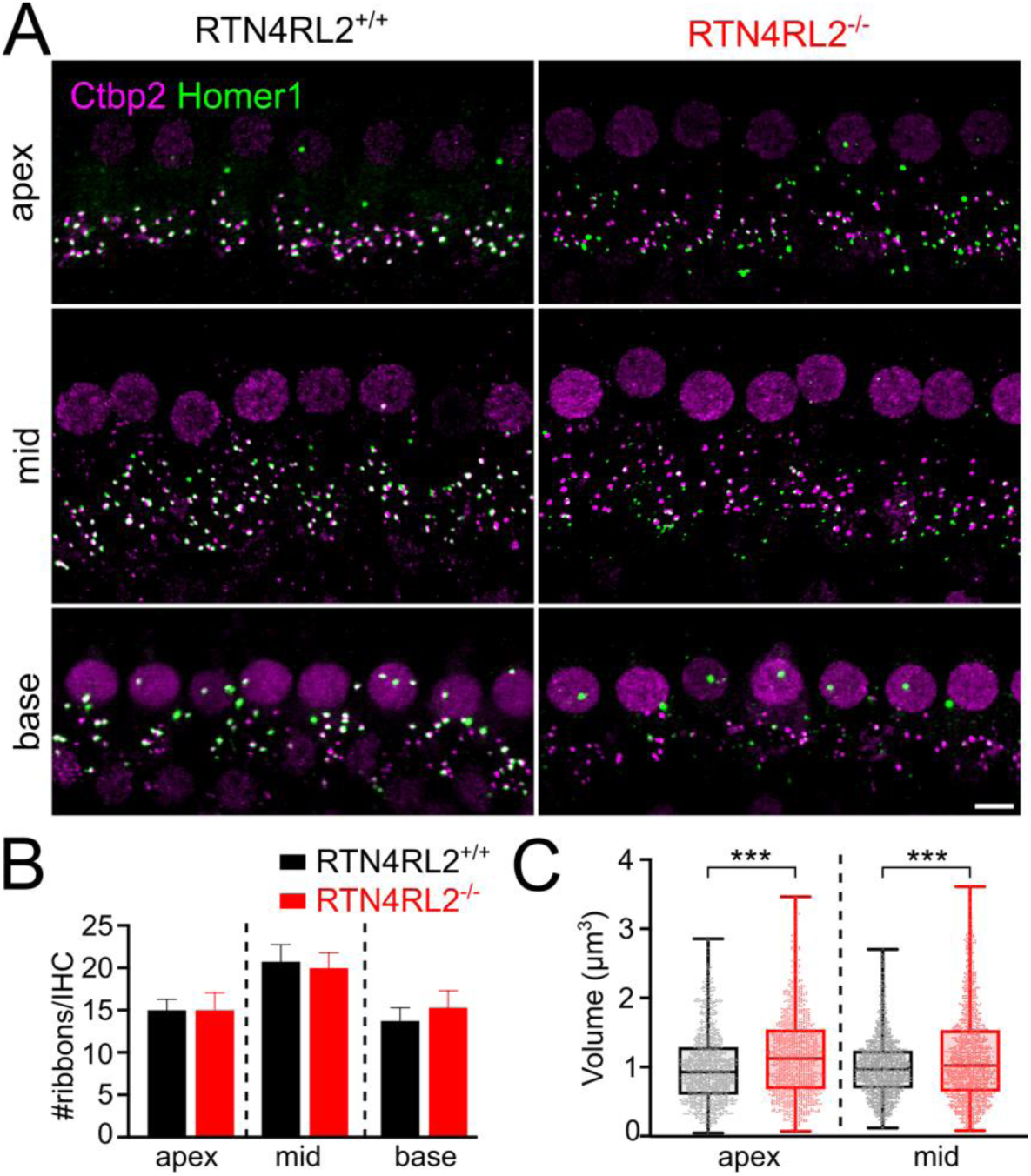
Intact number but enlarged size of the ribbons in RTN4RL2^-/-^ IHCs. (**A**) Maximum intensity projections of representative IHCs from apical, mid and basal regions of RTN4RL2^+/+^ (left) and RTN4RL2^-/-^ (right) cochleae of p21-30 mice. Synapses are visualized by staining against Ctbp2/Ribeye (ribbons) and Homer1 (PSDs). Scale bar = 5 μm. (B) The number of the ribbons is not affected along the tonotopic axis in RTN4RL2^-/-^ cochleae. (C) The size of the ribbons is increased in IHCs of both apical and middle turns in RTN4RL2^-/-^ cochleae (p < 0.001, Mann-Whitney-Wilcoxon test). N = 6 animals/genotype. Box-whisker plots show the median, 25/75 percentiles (box) and the range (whiskers). Individual data points are overlaid.

**Figure 2-figure supplement 3.**
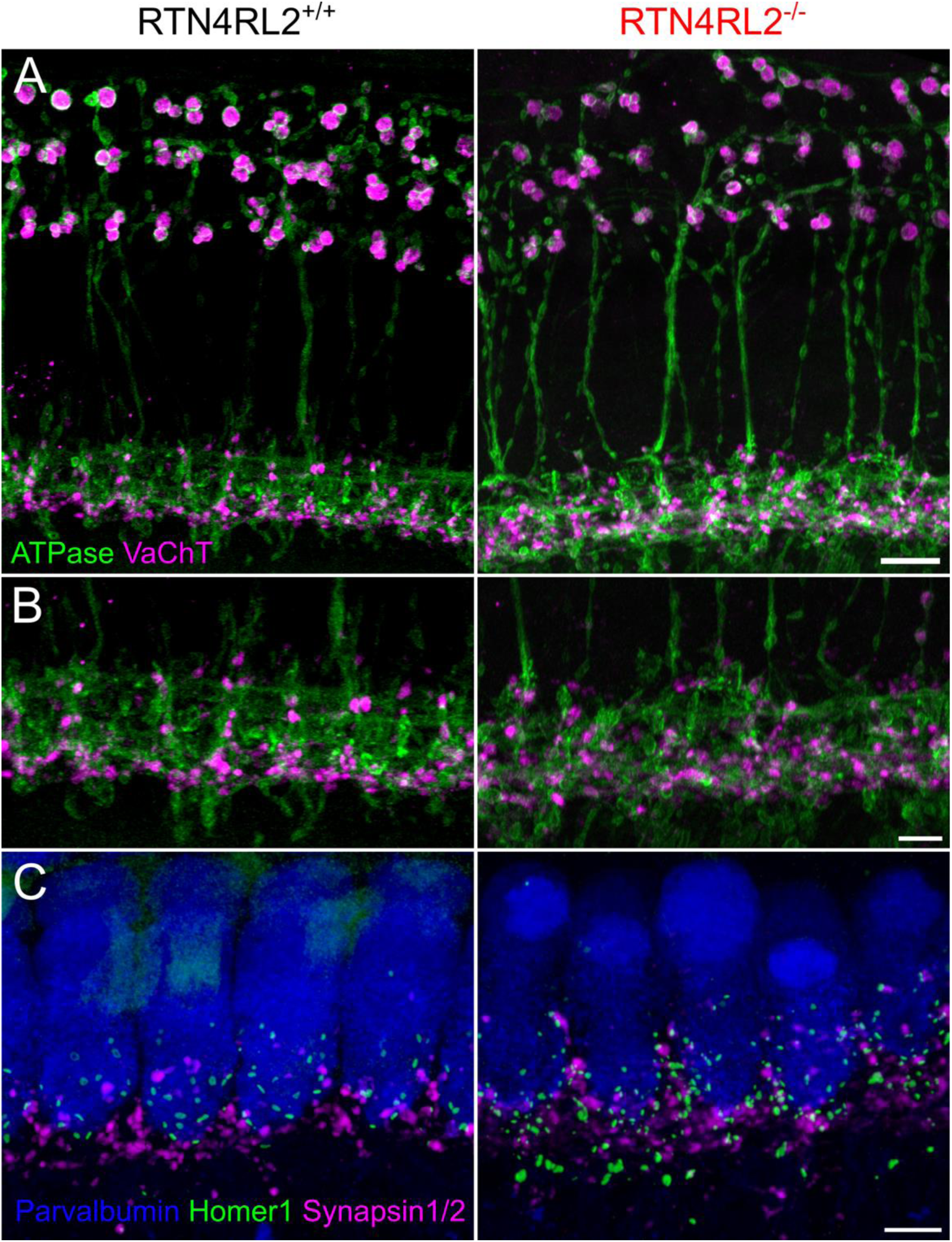
Efferent innervation pattern in RTN4RL2^-/-^ cochleae. (**A**) Maximum intensity projections of confocal stacks of outer and inner hair cell regions at the apical region of the cochlea from p21-30 RTN4RL2^+/+^ (left) and RTN4RL2^-/-^ (right) mice. SGN fibers/terminals and efferent terminals are visualized staining for Na^+^/K^+^ ATPase and vesicular acetylcholine transporter (VaChT), respectively. Scale Bar = 10 μm (**B**) Same as (A) but zoomed into the IHC region. Scale Bar = 5 μm (**C**) Maximum intensity projections of confocal stacks of a row of apical IHC region from p21 RTN4RL2^+/+^ (left) and RTN4RL2^-/-^ (right) mice. Presynaptic efferent terminals were stained with an anti-synapsin1/2 antibody. Scale bar = 5 μm.

**Figure 3-figure supplement 1.**
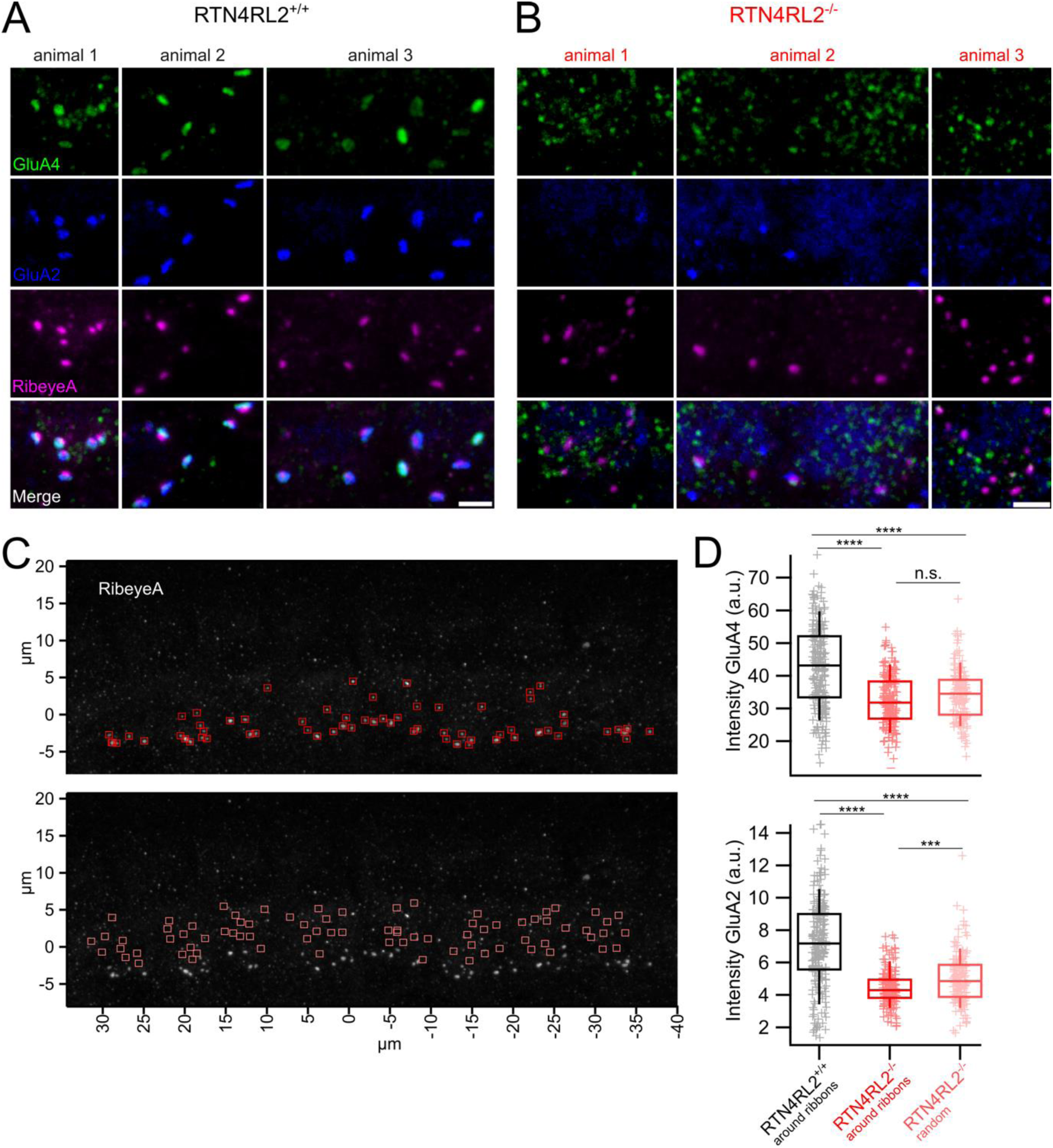
No apparent GluA4 signal juxtaposing presynaptic ribbons at IHC synapses of RTN4RL2^-/-^ mice. (**A, B**) Maximum intensity projections of a few imaging planes from apical IHC synaptic regions of 14-week-old N = 3 RTN4RL2^+/+^ (A) and N = 3 RTN4RL2^-/-^ (B) mice immunostained against GluA4, GluA2 and RibeyeA. Scale bars = 2 μm. (**C**) Fluorescence intensities of GluA4 and GluA2 channels were analyzed as the sum of all pixel intensities in approximately 1×1×1 μm³ volumes around the center of mass of RibeyeA immunofluorescent puncta (red boxes, top image). Background GluA4 and GluA2 fluorescence signals were calculated in the same way but outside the ribbons, at random spots inside IHCs (pink boxes, bottom image). Images show the maximum intensity projection of the IHC region in an RTN4RL2^-/-^ mouse. (**D**) Fluorescence intensities of GluA4 (top) and GluA2 (bottom) channels at ribbons of RTN4RL2^-/-^ mice do not show a significant increase compared to the background (RTN4RL2^-/-^ random) but are significantly decreased compared to RTN4RL2^+/+^ (Mann-Whitney-Wilcoxon test followed by Bonferroni multiple-comparison correction). Box-whisker plots show the median, 25/75 percentiles (box) and 10/90 percentiles (whiskers). Individual data points are overlaid.

**Figure 4-figure supplement 1.**
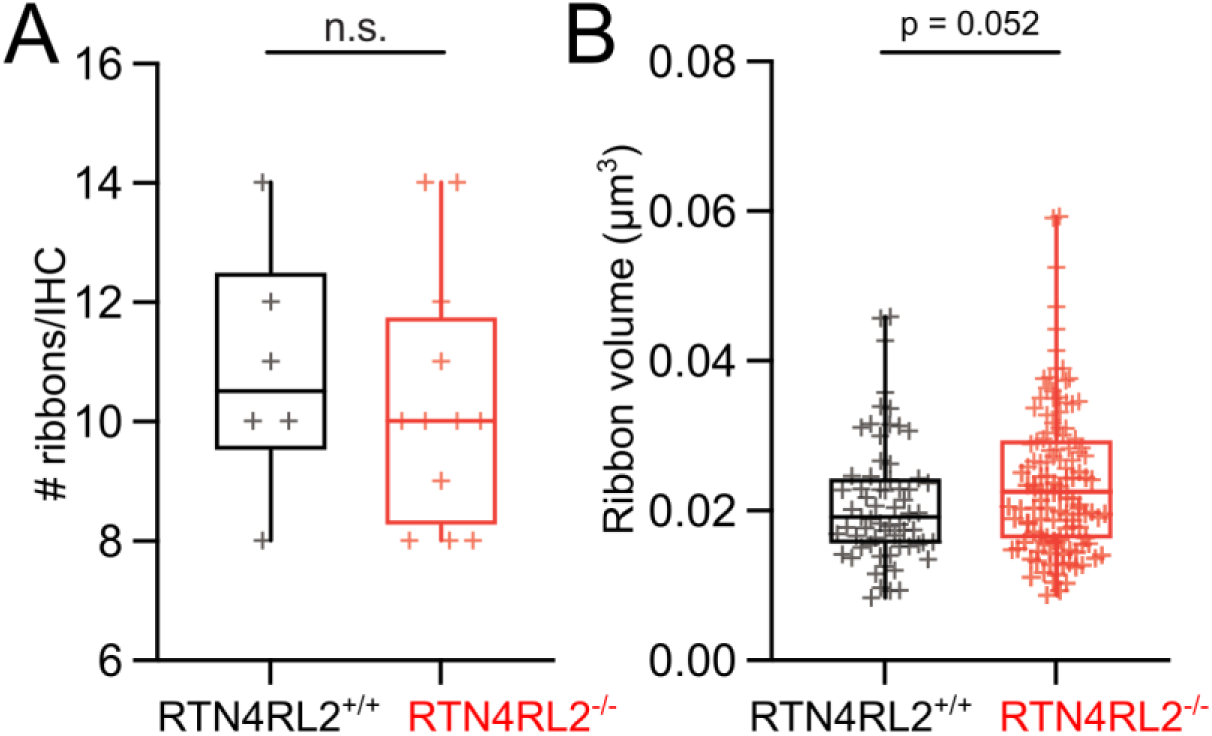
Quantification of ribbon number and volume in SBEM reconstructions. (**A**) The number of ribbons is not changed in RTN4RL2^-/-^ IHCs (RTN4RL2^-/-^: 10.3 ± 0.61, SD = 2.1, n = 12, N = 2 vs. RTN4RL2^+/+^: 10.8 ± 0.83, SD = 2.04, n = 6, N = 1; p = 0.54, Mann-Whitney-Wilcoxon test). (**B**) Ribbon volumes tend to be larger in RTN4RL2^-/-^ IHCs without reaching statistical significance (RTN4RL2^-/-^: 0.024 ± 0.001 μm^3^, SD = 0.01 μm^3^, n = 125 ribbons in 12 IHCs, N = 2 vs. RTN4RL2^+/+^: 0.021 ± 0.001 μm^3^, SD = 0.008 μm^3^, n = 65 ribbons in 6 IHCs, N = 1; p = 0.06, Mann-Whitney-Wilcoxon test). Box-whisker plots show the median, 25/75 percentiles (box) and the range (whiskers). Individual data points are overlaid.

